# Computational modelling of natural cell-to-cell heterogeneity reveals key parameters that control the diversity of human pancreatic islet β-cell excitability in response to glucose

**DOI:** 10.64898/2026.02.27.708039

**Authors:** I Goswami, J Koepke, M Baghelani, PE MacDonald, V Kravets, PE Light, AG Edwards

## Abstract

Insulin-producing β-cells demonstrate remarkable heterogeneity in their individual responsiveness to glucose, and that cellular heterogeneity contributes to coordinating islet activity and glucose homeostasis. Our current understanding of how variation in cell-intrinsic factors control cellular excitability and insulin secretion is informed by foundational experiments conducted on dispersed single β-cells. Such studies are limited in their ability to link multiple electrical or metabolic properties within a single cell and preclude the ability to relate, *post hoc*, each parameter’s contribution to glucose responsiveness. Computational modelling represents a unique and underutilized tool to integrate and investigate the role of natural β-cell heterogeneity in physiologic glucose responses. Herein, we utilize a high-volume single-cell electrophysiology “patch-seq” dataset to define the physiologically relevant sources of variability in human β-cell electrophysiology and model their influence on single-cell glucose responses. Three thousand *in silico* human β-cells were fitted to physiologically relevant variations in glucokinase activity, K^+^ current, Na^+^ current, Ca^2+^ current, and exocytotic function. Four dominant electrical phenotypes arose at low (2 mM) and high (20 mM) glucose: silent, bursting, spiking, and depolarized. Approximately 50% of uncoupled β-cells remained electrically silent at high glucose. Furthermore, Na^+^ channel half-inactivation voltage was a major predictor of the silent and spiking phenotypes at each glucose concentration, and of cells that transition from silent to spiking when glucose increased. Indeed, experimentally observed variation in Na^+^ channel voltage dependence was second only to variation in ATP-sensitive potassium channel conductance in determining β-cell excitability. Our data-driven computational modelling highlights the functional importance of electrical heterogeneity in human β-cell glucose responses, and provides a useful tool for generating testable hypotheses.

## INTRODUCTION

Pancreatic islets are networks of endocrine cells, each with a distinct role in maintaining glucose homeostasis. Within the islet, β-cells secrete insulin in response to elevated plasma glucose. This secretory process is coordinated, in large part, through cell-intrinsic electrical properties, linking nutrient metabolism and paracrine signalling to cellular excitability.^1,2^ The result is a tightly regulated system capable of maintaining stable blood glucose concentrations under varying physiological conditions. A growing body of evidence suggests that β-cell heterogeneity is a key determinant in coordinating insulin release, with differences in glucose transport and metabolism, ion currents, and Ca^2+^ dynamics contributing to their individual glucose responsiveness.^1–4^ These cells display broad variability in glucose responsiveness, ion channel expression, and hormone secretion, which together contribute to the dynamic range of islet hormone release and adaptability under physiological stress.

Despite recognizing cellular heterogeneity as functionally important to β-cell excitability, identifying how variability in specific metabolic or electrophysiological parameters gives rise to population-level coordination remains a major challenge. Experimental techniques such as live-cell Ca^2+^ imaging and optogenetic stimulation have been instrumental in characterizing cell-to-cell differences in excitability within intact islets, where electrical and paracrine coupling are preserved.^1,5–12^ However, these methods are limited in their ability to perturb individual parameters in isolation. For instance, altering the conductance of a single ion channel or transporter often introduces secondary effects that obscure the direct contribution of the targeted component. Moreover, the interdependence of metabolic and electrophysiological processes within β-cells complicates efforts to experimentally dissect the mechanisms that govern glucose responsiveness. As a result, the relative importance of specific electro-metabolic parameters for β-cell excitability and islet coordination remains incompletely understood and cannot be deconvolved with existing *in vitro* methodology.

Mathematical modelling allows systemic exploration of β-cell properties and their relative contributions to glucose responsiveness.^6,10–18^ By integrating experimental data into quantitative frameworks, computational modelling enables systematic exploration of the relationships between molecular components. Population-of-models (PoM) frameworks extend this concept by simulating ensembles of virtual cells, each defined by a unique combination of parameters drawn from experimentally constrained distributions. We extend the PoM approach, initially developed for cardiac electrophysiology studies, to capture β-cell heterogeneity in ion conductances and voltage dependencies, as well as metabolic fluxes.^19–31^ This strategy allows us to reproduce both the average and distributed properties observed in voltage-clamped primary cells, thereby providing a platform for testing how variation in specific parameters influences collective output (glucose-induced excitability) while preserving physiological realism.

While previous studies have relied on relatively small samples of dispersed β-cells to implement *in silico* heterogeneity,^32–35^ recent efforts have been made to generate larger datasets characterizing hundreds of primary human β-cells.^36^ Datasets of this scale permit a new approach to modelling heterogeneity in which variation in many components of the β-cell glucose response can be simultaneously varied and fitted to the measured distributions of electrophysiologic phenotypes. This allows the resulting models to capture a broad range of variation and covariation among molecular components while still constraining the behaviour of the collection of models to the measured function. Unlike earlier mathematical models based on rodent data or averaged human recordings, PoM-based frameworks can incorporate realistic ranges of conductance and voltage-dependence values to generate physiologically meaningful ensembles of β-cells.^32,37,38^ This approach enables rigorous testing of hypotheses about how electrometabolic variability influences glucose-stimulated electrical responses and insulin secretion.

In the present study, we implement this PoM approach to investigate the functional consequences of sodium current (*I*_*Na*_) heterogeneity in human β-cells. *I*_*Na*_ contributes less to β-cell excitability in mice because murine β-cell *I*_*Na*_ exhibits a very negative voltage dependence, leaving many channels inactivated at physiologic membrane potentials.^37^ In contrast, human β-cells display a bimodal distribution of *I*_*Na*_ voltage-dependence, and substantial but variable *I*_*Na*_, suggesting species-specific roles in depolarization and spike generation.^32,37,38^ Using a large dataset from Camunas-Soler *et al*.,^36^ we generated a population of β-cell models parameterized by experimentally derived conductances and voltage dependences for *I*_*Na*_, *I*_*KATP*_, *I*_*BK*_, and glucokinase activity.^32–35^ To capture the coupling between metabolism and electrical activity, we modulated an ATP-mimetic variable that links glycolytic flux, ATP consumption, and membrane excitability extrapolated from previous models.^39^ This framework allowed us to determine whether physiological variability in *I*_*Na*_ voltage dependence is sufficient to predict the emergence of excitable versus quiescent phenotypes across a population of human β-cells.

By simulating β-cell variability within an islet-focused modelling framework, this study contributes to a growing effort to understand how molecular diversity gives rise to coordinated islet function. Integrating experimental datasets with population-based computational models enables the dissection of mechanisms that cannot be isolated *in vitro*, providing a foundation for future work linking single-cell properties to multicellular behaviour in islet health and in states of metabolic impairment like types 1 and 2 diabetes.

## RESULTS

### Validation of in silico β-cell population heterogeneity

Our final population incorporated 16 heterogeneous parameters (Table 1) that faithfully recapitulated the variability observed in primary human β-cells (Figure 1). We validated the heterogeneity of our optimized *in silico* β-cell population by simulating parameter-specific voltage protocols and comparing the lognormal distribution of model parameters with primary data from Camunas-Soler *et al*. These simulations include steady-state inactivation (Figure 1A), total current activation (Figure 1D), and capacitance pulse-train protocols (Figure 1G), which are used to generate representative traces of the minimum, median, and maximum time series for key electrophysiologic measurements.

**Table 1.** Heterogeneous parameters in the population. The coefficient of variation (*σ*) was applied as a normalized form of parameter variability to construct the population of β-cell models. For parameters *V*_*GK,max*_ through *ḡ*_*KATP*_ (parameters 1-6), *σ* values were taken directly from the source datasets. The remaining 11 parameters were fit to the voltage-clamp distributions in Camunas-Soler *et al*. 2020 via differential evolution optimization.^36^ Initial guesses for these fitted *σ* values were taken from Braun *et al*. 2008,^32^ Takeda *et al*. 1993,^70^ Gidh-Jain *et al*. 1993,^71^ and Speier *et al*. 2005.^72^ The optimization was constrained to allow *σ’s to vary from 0*.*5 to 2-fold of the initial value for all conductance parameters, and 0*.*33 to 3-fold for all voltage-dependence parameters. The fraction, ρ*_*high*,_ *was also optimized from an initial estimate of 0*.*5*.

| Parameter Number | Parameter Name | SD/mean ($\sigma$) | | Parameter Description | Source |
| --- | --- | --- | --- | --- | --- |
| | | Initial $\sigma$ | Fitted $\sigma$ | | |
| 1 | $V_{GK,max}$ | 0.710 | Not fitted | Maximum flux of GK | Takeda *et al.* 1993 |
| 2 | $K_{GK}$ | 0.900 | Not fitted | Glucose concentration at half-activation of GK | Gidh-Jain *et al.* 1993 |
| 3 | $g_{Kv}$ | 0.370 | Not fitted | $I_{Kv}$ maximal conductance | Braun *et al.* 2008 |
| 4 | $g_{BK}$ | 0.800 | Not fitted | $I_{BK}$ maximal conductance | Braun *et al.* 2008 |
| 5 | $g_{KATP}$ | 0.940 | Not fitted | $I_{KATP}$ maximal conductance | Speier *et al.* 2005 |
| 6 | $g_{CaL}$ | 0.220 | 0.206 | $I_{CaL}$ maximal conductance | Camunas-Soler *et al.* 2020 |
| 7 | $g_{CaPQ}$ | 0.590 | 1.180 | $I_{CaPQ}$ maximal conductance | Camunas-Soler *et al.* 2020 |
| 8 | $g_{CaT}$ | 0.400 | 0.483 | $I_{CaT}$ maximal conductance | Braun *et al.* 2008 |
| 9 | $g_{Na,high}$ | 0.490 | 0.671 | $I_{Na,high}$ maximal conductance | Camunas-Soler *et al.* 2020 |
| 10 | $V_{hNa,high}$ | 0.140 | 0.175 | $I_{Na,high}$ half inactivation potential | Camunas-Soler *et al.* 2020 |
| 11 | $V_{mNa,high}$ | 0.140 | 0.175 | $I_{Na,high}$ half activation potential | Camunas-Soler *et al.* 2020 |
| 12 | $n_{hNa,high}$ | 0.15 | 0.263 | $I_{Na,high}$ inactivation slope factor | Braun *et al.* 2008 |
| 13 | $g_{Na,low}$ | 0.490 | 0.679 | $I_{Na,low}$ maximal conductance | Camunas-Soler *et al.* 2020 |
| 14 | $V_{hNa,low}$ | 0.140 | 0.067 | $I_{Na,low}$ half inactivation potential | Camunas-Soler *et al.* 2020 |
| 15 | $V_{mNa,low}$ | 0.140 | 0.067 | $I_{Na,low}$ half activation potential | Camunas-Soler *et al.* 2020 |
| 16 | $n_{hNa,low}$ | 0.15 | 0.101 | $I_{Na,low}$ inactivation slope factor | Braun *et al.* 2008 |
| Parameter Number | Parameter Name | Initial $\rho_{high}$ | Fitted $\rho_{high}$ | Parameter Description | Source |
| 17 | $\rho_{high}$ | 0.500 | 0.370 | Fraction of cells expressing $I_{Na,high}$ | no initial source |

**Figure 1.**
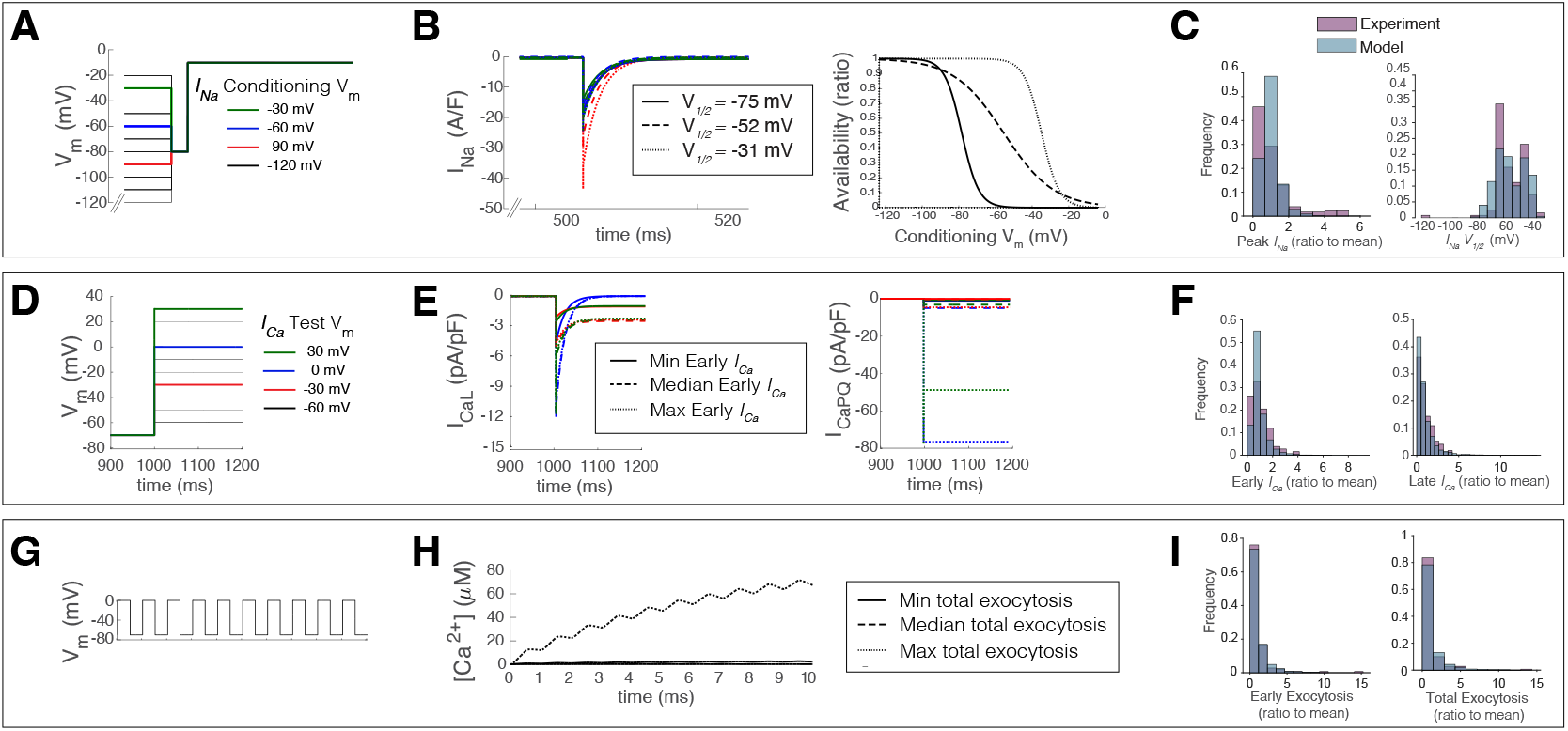
Variation in voltage clamp function of the heterogeneous β-cell population. (**A**) Steady-state inactivation voltage-clamp protocol with colour-coded voltage waveforms is used to generate representative traces in panel B and B’. (**B**) Representative INa time-series and Boltzmann fits for three cells spanning the range of steady-state inactivation voltage-dependence. (**C**) Histograms for the lognormal distribution of peak INa and the bimodal distribution of INa half-inactivation voltage in the modelled heterogeneous population (blue), and as measured by Camunas-Soler et al.^36^ (violet). (**D**) Total activation voltage-clamp protocol with colour-coded voltage waveforms, used to generate representative traces in panels E and E’. (**E**) Representative ICaL and ICaPQ for the three cells with maximum, minimum and median Early ICa. (**F**) Histograms for the lognormal distributions of Early and Late ICa for the model population (blue) and data from Camunas-Soler et al. ^36^ (violet). (**G**) A pulse-train voltage-clamp protocol was used to generate representative traces in panel H. (**H**) Representative cytosolic Ca^2+^ responses, which we us as a proxy for exocytosis responses indicated by the progressive increase in capacitance elicited by this protocol in the experimental dataset. (**I**) Histograms for the lognormal distributions of Early and Total exocytosis responses for the model population (blue) and exocytosis data from Camunas-Soler et al.^36^ (violet).

We plotted representative *I*_*Na*_ envelopes (Fig 1B) and Boltzmann fits (Fig 1B’) for cells with maximum (-31mv), median (-52mV), and minimum (-75mV) *I*_*Na*_ half-inactivation voltages. These data demonstrate the effect of voltage dependence on channel availability and peak I_Na_. Sigmoidal fits from our simulated population closely match log-normalized experimental data from Camunas-Soler *et al*., where we observed a similar left-skew in peak *I*_*Na*_ distribution without data-trimming (Fig 1C) and bimodal distribution of half-inactivation voltages (Fig 1C’). In our optimized population, 37% of cells were sampled from the *I*_*Na,high*_ and 63% from the *I*_*Na,low*_ half-inactivation peaks (ρ = 0.37) to achieve physiologically relevant bimodality.

We also generated representative *I*_*CaL*_ and *I*_*CaPQ*_ traces (Fig 1E, E’), the two dominant contributing ion currents in the total-activation protocol (Fig 1D). Once again, the lognormal distribution of early and late *I*_*Ca*_ components was similar to primary data and exhibited a pronounced left-skew (Fig 1F). Finally, we simulated intracellular Ca^2+^ responses to the pulse-train protocol, as a proxy for exocytosis (Fig 1H). The lognormal distribution of simulated early and late exocytosis aligned with the Camunas-Soler *et al*. dataset (Fig 1I). Having captured appropriate variance in ion channel kinetics and glucokinase activity, we proceeded to implement physiologically realistic electro-metabolic coupling.

### Glucose-responsiveness and optimization of electro-metabolic coupling

Our optimized *in silico* population of 3,000 β-cells exhibited four distinct electrophysiological phenotypes: (1) electrical silence (complete silence or sub-threshold depolarization), (2) bursting, (3) spiking, and (4) a small number (< 10 cells) became permanently depolarized with no oscillatory phenotype in at least one of the two glucose concentrations (Fig 2A). These four behaviors are coupled to metabolic activity in the model, and when stimulated by glucose, the metabolic subsystem oscillates on a seconds-to-minutes timescale. However, the coupling between metabolic activity and electrical activity is variable in the model population, as it is in primary β-cells. That is, individual models can be electrically active (spiking, bursting) in the absence of metabolic activity, and metabolically active cells can remain electrically silent if electrometabolic coupling is weak. To characterize this relationship in the model population we used the ATP-mimetic variable, *a* (Equation 1), as a proxy to measure metabolic activity.Specifically, if *a* exceeded a threshold of 2.0 mM, the cell was deemed to be metabolically oscillating and active (Fig 2B).

**Figure 2.**
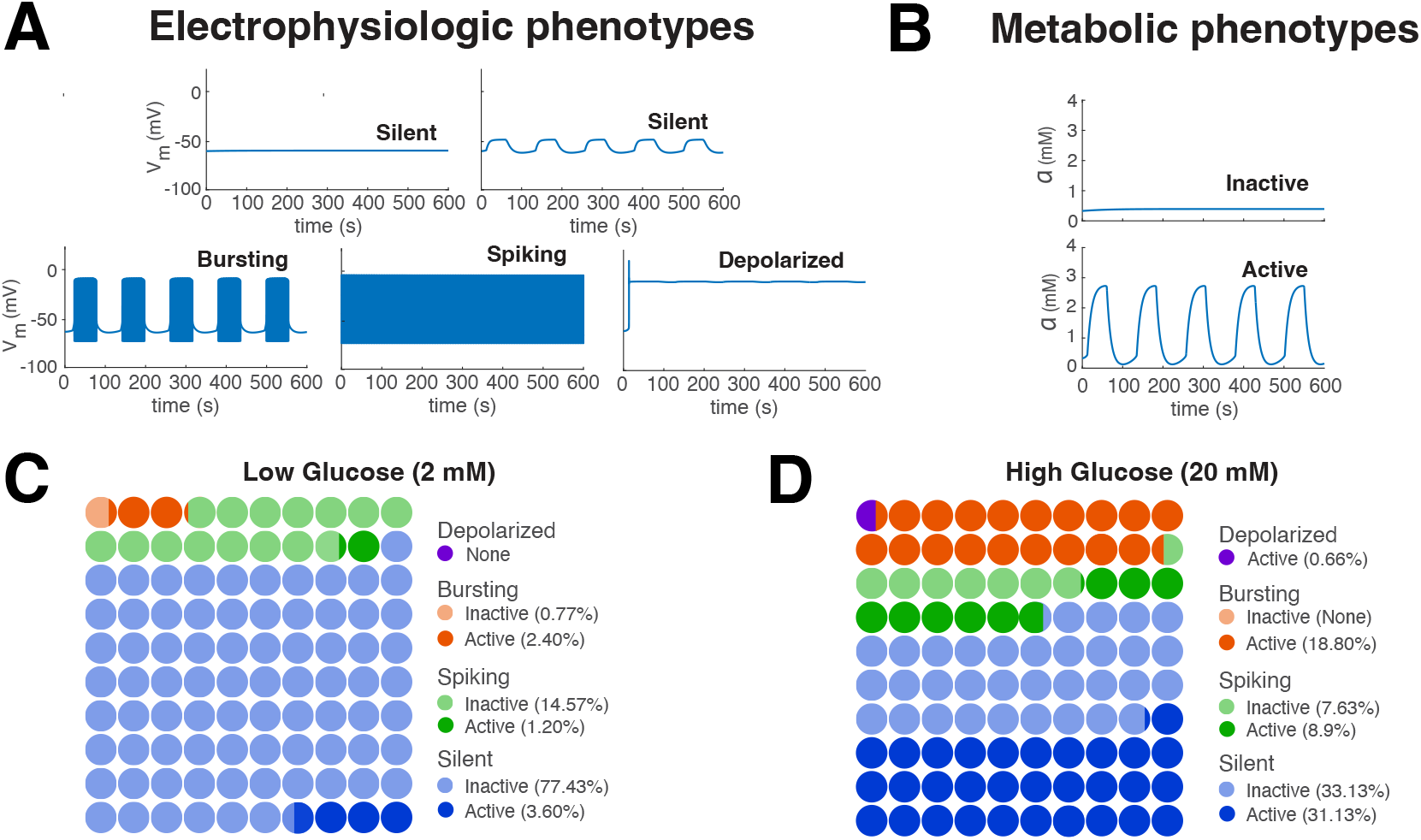
Glucose response classes of the optimized β-cell population. **(A)** Physiologic classes of electrical activity (Silent, Bursting, Spiking and Depolarized) were determined from membrane potential (Vm) time series at 2 and 20 mM glucose. Electrically silent cells commonly exhibited subthreshold depolarization at high glucose concentrations in response to metabolic activation e.g. top-right trace. Electrical activity was accompanied by either inactive or active metabolic states based on oscillatory dynamics of the ATP mimetic variable, a. (**C**) parts-of-whole representation of metabolic and electrical phenotypes at 2mM glucose. (**D**) parts-of-whole representation of metabolic and electrical phenotypes at 20mM glucose.

At low glucose, 81% of cells were electrically silent, while 19% exhibited electrical activity (16% spiking, 3% bursting). Despite nearly one-fifth of cells being electrically active, only 7.2% were metabolically active at 2 mM glucose, as determined by the ATP-mimetic threshold. Of the metabolically active cells, 108 were silent, 36 spiking, and 72 bursting. Most spiking cells were metabolically inactive at low glucose, while most bursting cells were metabolically active. Over 95% of electrically silent cells were also metabolically silent (Fig 2C).

At 20mM glucose, 59.2% of cells exhibited metabolic activity, and 58.8% were electrically active. The bursting phenotype accounted for 18.8% of cells, all of which were metabolically active. Spiking cells increased from 16% in low glucose to 16.5% of cells in high glucose conditions, with only 53% of actively spiking cells exhibiting metabolic activity. Approximately 48.4% of electrically silent cells were metabolically active in high glucose conditions (Fig 2D).

### Contribution of electro-metabolic heterogeneity to glucose-response phenotype

Having determined the baseline characteristics of our electro-metabolically optimized β-cells, we chose to investigate the contribution of heterogeneity in individual parameters to each electrical phenotype. We conducted both group-wise tests to assess mean effects and logistic regression to decompose the simultaneous variation across all 16 heterogeneous parameters. The coefficients returned by logistic regression indicate the sensitivity of each phenotype (as compared to all other phenotypes) to the parameter in question.

K_ATP_ conductance (ḡ_KATP_) was the strongest predictor of the silent and spiking phenotypes, where increased conductance favoured electrical silence and decreased conductance promoted spiking at both glucose concentrations. Glucokinase maximum reaction velocity (V_GK,max_) and dissociation constant (K_GK_) were unique and sensitive predictors of the bursting phenotype where high V_GK,max_ and low K_GK_ favoured bursting at both glucose concentrations. The large conductance Ca^2+^-activated K^+^ current (BK) was implicated in all three electrical phenotypes, although its role in electrical bursting was most pronounced. Because the Spiking and Silent phenotypes exhibit equal and opposite directions of ḡ_KATP_ sensitivity, their combination in the logistic model is indistinguishable from bursting. This is confirmed by the ANOVA analyses (Figure 3A) and the underlying ḡ_KATP_ distributions for Bursting and non-bursting cells (Figure 4C), which indicate that Silent and Bursting cells have similar mean ḡ_KATP_, and the Bursting distribution is spanned by non-bursting cells.

**Figure 3.**
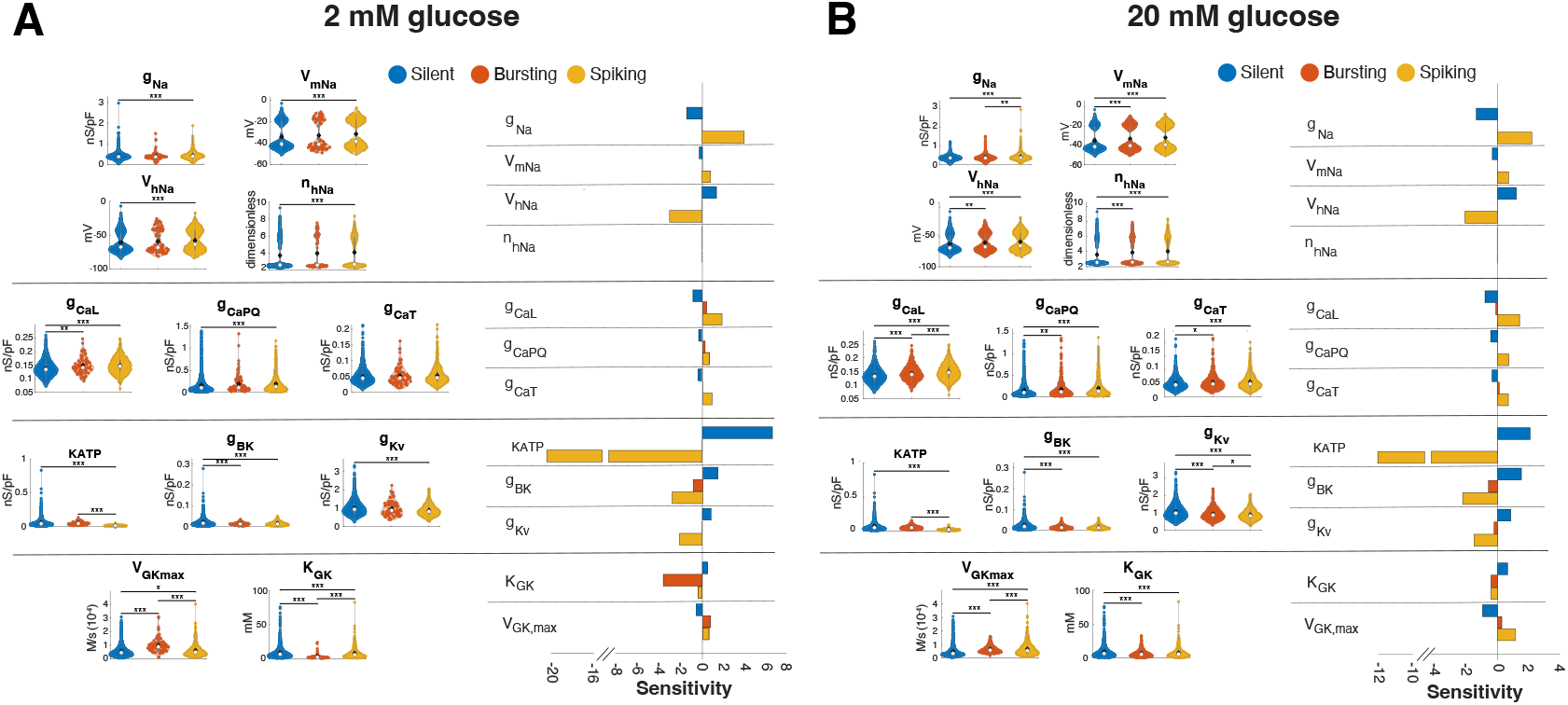
I_Na_ heterogeneity contributes differentiating Silent and Spiking cells at 2 mM and 20 mM glucose. (Both Panels). Left side presents violin plots of parameter values for cells in each of the 3 major phenotypes (Silent, Bursting and Spiking). Mean (white circles) and median (black diamonds) are provided with the full distributions. Right panels show all significant coefficients (sensitivities) for logistic regression performed on the standardized parameter deviations for the 3 phenotypes. Non-significant coefficients are presented as 0. **(A)** At low glucose, I_KATP_ parameters (ḡ_KATP_) and I_Na_ parameters are key for differentiating the Silent and Spiking phenotypes, while the GK parameters K_GK_, and V_GK,max_ are unique in selectively distinguishing Bursting from the other two phenotypes. (**B**) Similar patterns of effect at high glucose, with generally larger effect sizes than at 2 mM, and notably enhanced sensitivity of Bursting to the large conductance Ca^2+^-activated K^+^ current (g_BK_ parameter).

**Figure 4.**
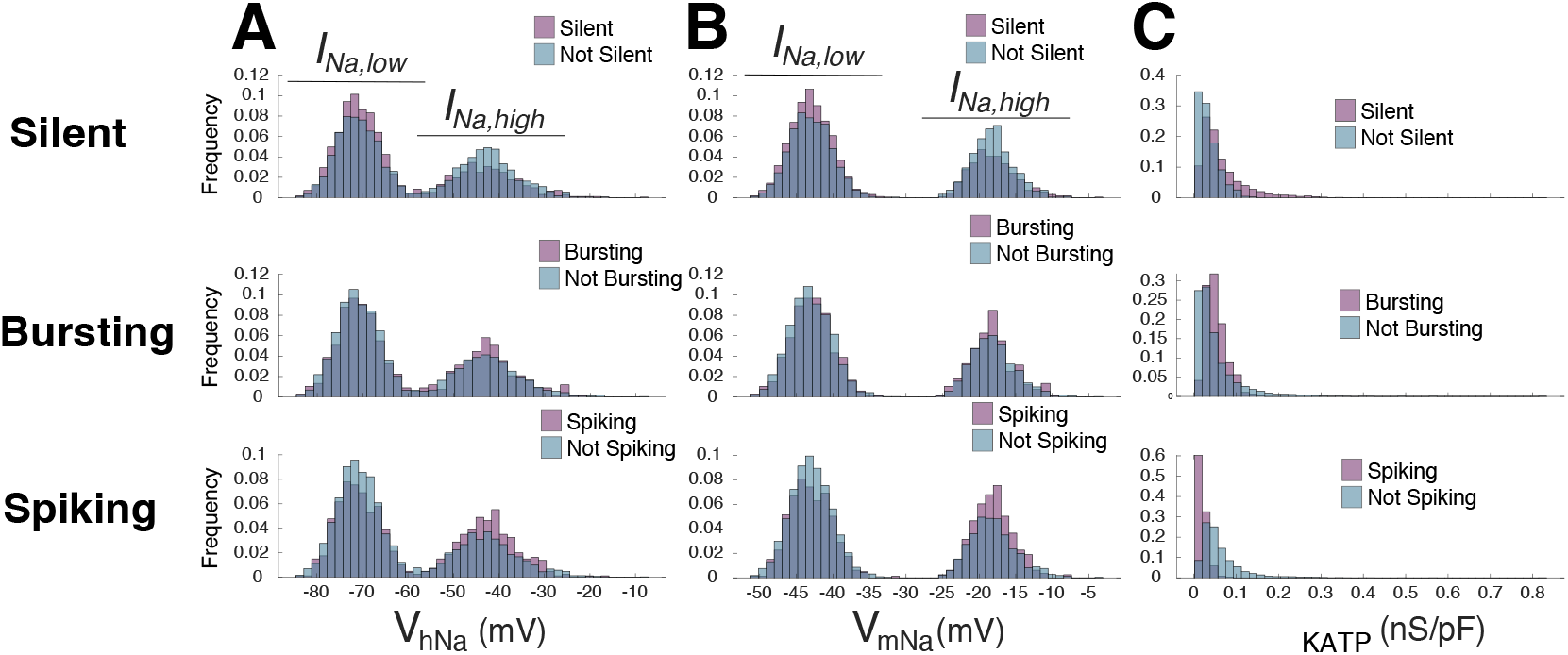
The sensitivity of I_Na_ voltage-dependence is defined by a shifting balance between I_Na,low_ and I_Na,high_. Distributions for the I_Na_ half-inactivation parameter, V_hNa_ **(A)**, the I_Na_ half-activation parameter V_mNa_ **(B)**, and maximal I_KATP_ conductance, ḡ_KATP_ **(C).** All three parameters are shown for the 3 main phenotypes under high glucose conditions. Both the V_hNa_ and V_mNa_ distributions show reciprocal enrichment of the left-shifted I_Na,low_ component of the distribution for Silent cells, and the right-shifted I_Na,high_ component for Spiking cells. Bursting cells are not clearly differentiated from non-Bursting cells by these two parameters. Similarly, higher ḡ_KATP_ is enriched among Silent cells and vice versa for Spiking cells. Bursting is again intermediate and indistinguishable from the combination of the other two classes.

Inactivation voltage dependence of *I*_*Na*_ (V_hNa_) and Na channel conductance (g_Na_) are among the strongest predictors of spiking and silent β-cells, irrespective of simulated glucose level. V_hNa_ is a positive predictor of the silent phenotype, where a left-shifted, more negative, half-inactivation voltage promotes electrically silent cells at both glucose concentrations. The same effects exist between low g_Na_ and silent cells. The reciprocal is true for both parameters in spiking cells, where more positive V_hNa_ and high g_Na_ increase the likelihood of electrical spiking (Fig 3A, 3B).

Since we varied the three voltage dependence parameters in coordination to retain physiologic steady-state properties (half inactivation/V_hNa_, half activation/V_mNa_, and slope of half inactivation), it was not possible to distinguish which parameter was most important for the observed sensitivity. To address this, we plotted their distributions to visualize the contribution of voltage-dependence parameters to each electrical phenotype (Fig 4A, 4B). These show a greater proportion of right-shifted voltage-dependence for cells in the spiking class, and left-shifted for silent. Because V_hNa_ and V_mNa_ were varied simultaneously, the underlying mechanism reflects the combined effect of both changes in voltage dependence.

### Heterogeneity dictates the transitional phenotype from low to high glucose

Having ranked the importance of model parameters to electrical phenotypes in either glucose condition, we aimed to understand the factors contributing to glucose excitability. That is, the change in electrical phenotype elicited by increasing glucose. We applied the same approaches to identify a hierarchy of parameters in the different glucose-dependent transitional phenotypes. Given the large population of electrically silent cells at low glucose, this analysis only considered the silent-to-silent (Si-Si), silent-to-bursting (Si-Bu), or silent-to-spiking (Si-Sp) phenotypes. Si-Si cells accounted for 65% of cells, while Si-Bu and Si-Sp comprised 27% and 7% of the remaining population, respectively. 1% of cells adopted an uncommon bursting-to-spiking phenotype, although we excluded these from subsequent analyses.

Left-shifted *I*_*Na*_ voltage dependence was among the more sensitive predictors of the Si-Si subset (Figure 5B, top-right), along with gBK, and ḡ_KATP_. The sensitivities are reciprocal for Si-Si and both Si-Bu and Si-Sp for all parameters, which reflects the general influence of each parameter on glucose-induced cellular excitability. Surprisingly, V_GK,max_ is not a sensitive predictor of Si-Bu, although this is likely explained by the fact that the Si-Bu distribution for V_GK,max_ is completely spanned by both the Si-Si and Si-Sp distributions (Figure 5B, bottom-left). This blunts the ability of the dichotomous logistic analysis from identifying variation in V_GK,max_ as distinguishing Si-Bu cells from the combination of all Si-Si and Si-Sp cells. Since the mean differences in V_GK,max,_ among the three transition classes were large, it is likely that the logistic sensitivity does not fully reflect the importance of VGK,max.

**Figure 5.**
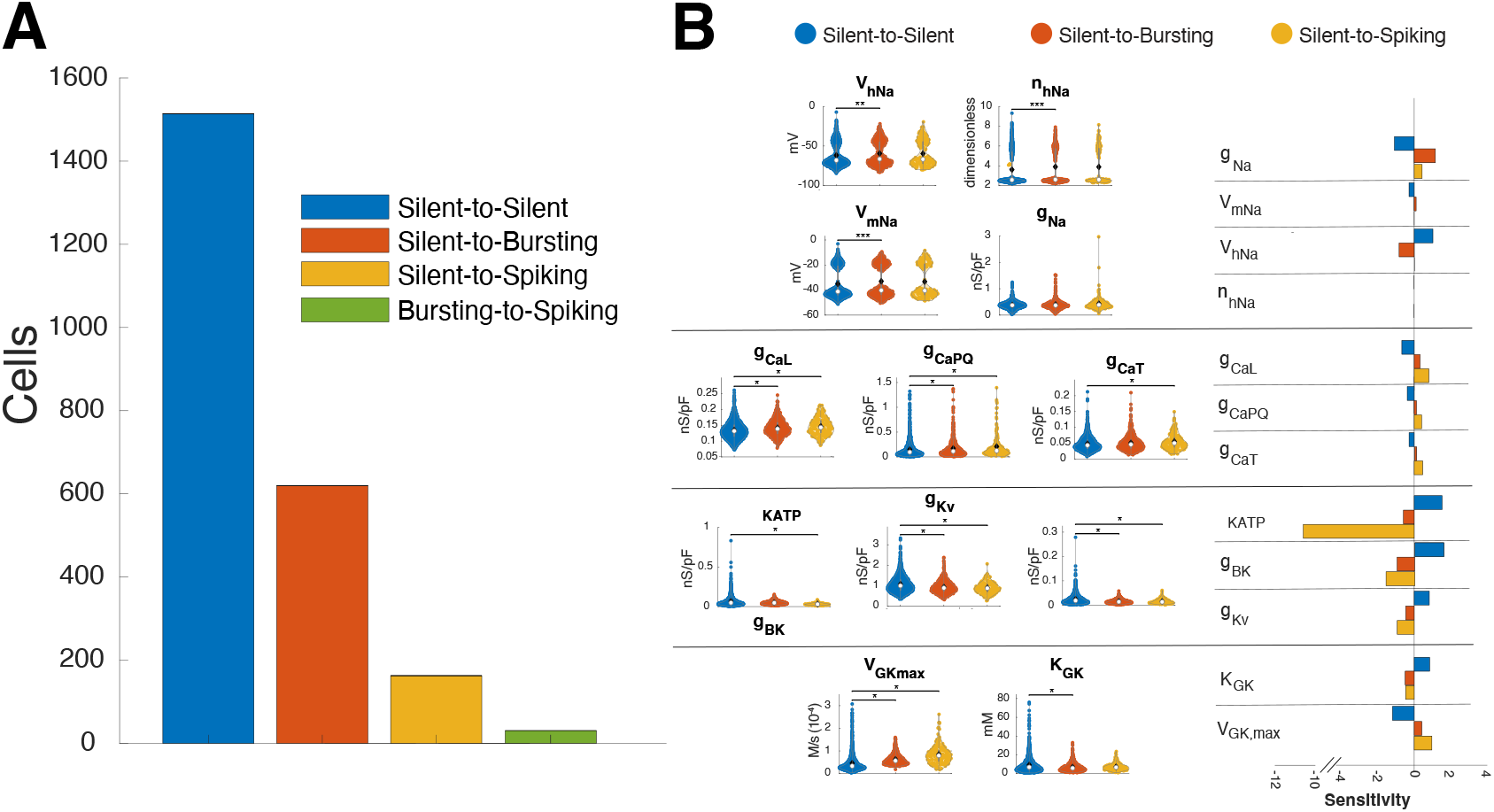
Left-shifted INa voltage-dependence stabilizes the Silent phenotype during glucose-stimulation, and right-shifted voltage dependence may contribute to gucose-induced Bursting. **(A)** The number of cells for each glucose-induced phenotype transition. Most cells remained Silent after exposure to 20 mM glucose, but 26.7% transitioned to Bursting, and 7% to Spiking. **(B)** Left-shifted I_Na_ voltage-dependence and reduced g_Na_ were predictive of cells that remained Silent at both low and high glucose, although the mean effects were very subtle. Because Silentto-Bursting was the other major group the reciprocal sensitivities were observed for that group.

The Si-Sp group had a very similar parameter hierarchy as the group of cells that were Spiking at both low and high glucose (Figure 3). ḡ_KATP_ is the most sensitive determinant in all three of those subsets (Si-Sp, low glucose spikers, high glucose spikers), and supports the canonical interpretation that *I*_*KATP*_ plays a key role in maintaining stability close to resting potential. Interestingly, *I*_*Na*_ voltage dependence was not a significant predictive component for the 163 Si-Sp cells, even though it was predictive in the 611 cells that were Spiking at low glucose and 796 cells that were Spiking at high glucose. This emphasizes the importance of *I*_*Na*_ for setting the basal excitability of the β-cell population rather than for transducing metabolic activity into electrical activity.

### Modulating voltage dependence

To test the relative importance of *I*_*Na*_ voltage-dependence in glucose responsiveness, we modelled an entire population of cells with *I*_*Na,high*_ (*ρ* = 1.00) or *I*_*Na,low*_ (*ρ* = 0.00) distribution. At *ρ* = 1.00, the population exhibited a higher number of spiking cells at both basal and high glucose than at *ρ* = 0.00 (Figure 6B). The effects on glucose-induced bursting were subtle, where shifting the population from *ρ* = 0.00 to *ρ* = 1.00 only increased the number of silent-to-bursting cells by 8%. These data suggest that *I*_*Na*_ voltage dependence plays a crucial role in determining basal excitability but may not be sufficient to promote a significant expansion of the bursting class in response to glucose stimulation. *I*_*Na*_ heterogeneity contributed similarly to electrical excitability in both the optimized *ρ* = 0.37 and all-high *ρ* = 1.00 populations, albeit with lower sensitivity to other *I*_*Na*_ parameters in the all-high population. Notably, *I*_*Na*_ ceased to contribute to statistically discernible phenotypes within the *I*_*Na,low*_ (*ρ* = 0.00) β-cell population (Figure 6A).

**Figure 6.**
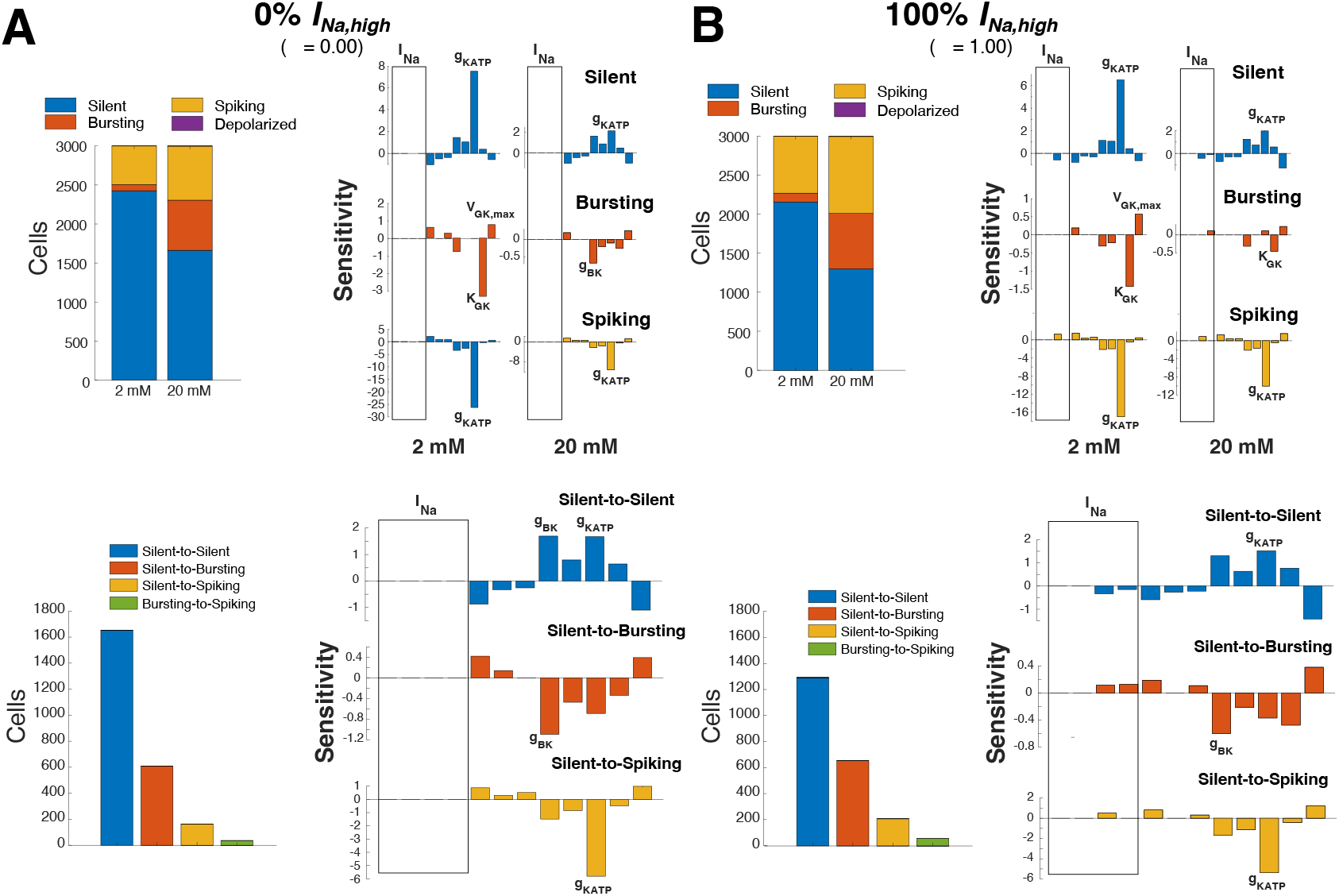
Bi-modal I_Na_ voltage-dependence stabilizes the Silent phenotype during glucose-stimulation, and right-shifted voltage dependence may contribute to glucose-induced Bursting. **(A)** The number of cells for each glucose-induced phenotype transition. Most cells remained Silent after exposure to 20 mM glucose, but 26.7% transitioned to Bursting, and 7% to Spiking. **(B)** Left-shifted I_Na_ voltage-dependence and reduced g_Na_ were predictive of cells that remained Silent at both low and high glucose, although the mean effects were very subtle. Because Silent-to-Bursting was the other major group the reciprocal sensitivities were observed for that group.

Together, these observations suggest that the bimodal distribution of *I*_*Na*_ voltage dependence properties may represent a key intrinsic mechanism by which β-cells tune their basal excitability within the islet.

## DISCUSSION

Here, we constructed a heterogeneous population of simulated human β-cells using a hybrid PoM approach. Constrained by high-quality primary electrophysiological datasets, our model offers a platform to investigate the implications of physiologically relevant β-cell electro-metabolic heterogeneity to electrical activity and glucose responsiveness. Such simulations have allowed us to characterize the hierarchy of parameters, such as *I*_*Na*_ voltage dependence and glycolytic flux on β-cell excitability and explore the effects of bimodal parameter distribution on β-cell function.

Our simulations accurately replicated observed distributions in voltage-clamp kinetics, β-cell basal activity, and population glucose responsiveness. Notably, we demonstrate that bimodal *I*_*Na*_ voltage dependence plays a defining role in basal excitability. We also find that robust parameter inference and investigation, enabled by large-scale modelling (n=3,000), is essential for examining subtle determinants of β-cell behaviour that would be undetectable in conventional computational studies (n<30). This novel approach has the potential to allow researchers to evaluate how changes in the activity and/or expression of electrometabolic machinery may impact the excitability of individual β-cells. Furthermore, this approach may be incorporated into drug discovery pipelines to screen the effects of ion channel modulators on predicted β-cell excitability.

### Bimodal I_Na_ voltage dependence

While prior studies have acknowledged differences in β-cells’ *I*_*Na*_ voltage-dependence, our PoM approach allowed us to explore the functional significance of this phenomenon. Past studies have characterized distinct *I*_*Na*_ voltage dependencies between human and murine β-cells, with mice exhibiting a pronounced left-shift that effectively silences *I*_*Na*_ under physiological conditions.^37,40,41^ In contrast, human β-cells express both left- and right-shifted *I*_*Na*_ with inactivation midpoints separated by ∼30mV.^36^ This key difference is likely attributed to the varying expression of NaV1.7 and NaV1.3 isoforms, which inactivate near -70mV and -40mV, respectively.^32,37,38^ While previous groups have hypothesized that a small population of cells with right-shifted *I*_*Na*_ voltage dependence could disproportionately influence excitability in a functionally coupled β-cell network,^38^ our study is the first to quantify its population-level impact. Indeed, this heterogeneity in voltage dependence appears to be a key determinant in basal electrical activity and may play a role in dictating individual β-cell function when uncoupled, as in single-cell experiments.

By explicitly incorporating *I*_*Na*_ bimodality in our study, we demonstrated that the presence of *I*_*Na,high*_ markedly increased the likelihood of an active (spiking or bursting) electrical phenotype, with a preference for electrical spiking. Logistic sensitivity analysis and modulation of the ratio of high and low *I*_*Na*_ populations (*ρ*) support our conclusion that the balance between populations of *I*_*Na*_ is a tunable determinant of basal excitability in human β-cells. Although it remains to be validated whether this spiking preference reflects a specialized physiological function or a general increase in excitability, the relative uncoupling of the spiking phenotype from metabolic characteristics suggests a possible role for basal insulin secretion.

These findings raise the intriguing possibility that NaV isoform expression might dictate β-cells’ intrinsic electrical function, and shifts in isoform expression may contribute to dysregulated activity in pathophysiology. Such shifts in isoform expression may be difficult to monitor with transcriptomic techniques and will likely require further functional studies or proteomic examination of single β-cells. Studying the bimodality of *I*_*Na*_ voltage dependence in β-cells from type 2 diabetic patients may provide insight into the distribution of NaV isoforms *in situ* and indicate whether there are overt changes in channel expression or perhaps more subtle changes in channel activity mediated by post-translational modifications. Phosphorylation events may be vital in defining the voltage dependence of existing channels, where stress-induced changes in intracellular kinase activity could feasibly influence β-cell excitability without altering channel assembly. Studies of NaV1.7 document a mild rightward shift in activating voltage dependence after PKC activation.^42^ As increased PKC activation is well-documented in dyslipidemia and insulin resistance,^43^ changes in intracellular kinase activity may subtly modulate β-cell voltage dependence and basal insulin secretion. Bimodal *I*_*Na*_ voltage dependence may represent a tunable determinant of β-cell excitability, potentially linking molecular plasticity in NaV channel regulation to basal insulin secretion and the altered electrical behaviour characteristic of diabetic islets.

### Electro-metabolic coupling and glucose responsiveness

Our model highlights the relationship between glucose metabolism and electrical behaviour in human β-cells and provides insight into the relative contribution of GK activity and K_ATP_ conductance. While these parameters were dominant determinants of glucose-stimulated bursting, we found spontaneous spiking to be largely glucose-independent. Compared to previous rodent studies, human β-cells exhibited reduced electro-metabolic coupling. This is consistent with reports of blunted metabolic amplification in human β-cells with higher basal activity dispersed cells compared to intact islets,^1,2,44–47^ although its physiological impact must be assessed in future population studies. Our simulations found 7.2% of cells metabolically active at 2mM glucose – substantially lower than the 20% basal activity observed in rodents – though the fraction of active cells at 20mM glucose was similar (∼59%). ATP consumption kinetics, represented by the coefficient *kA*, was a major determinant of glucose-mediated excitability in our modelled β-cell population. We noted that increasing the hydrolysis rate constant from 3e^-5^ to 4e^-5^ increased the active fraction from 35% to 49% in high glucose and increased electro-metabolic coupling fidelity to 0.47. These findings suggest that ATP production, turnover, and downstream ion currents modulate electrical heterogeneity in ways that metabolic input alone cannot capture.

### Implications for β-cell function and insulin secretion within islets

The differing electrical phenotypes observed in our model (silent, spiking, bursting) recapitulate known experimental features of heterogeneous human β-cell electrical function and Ca^2+^ dynamics. Approximately 19% (Fig. 2) of our β-cell population was spontaneously active at lower glucose, consistent with reports that a small fraction of β-cells may mediate basal insulin secretion in sub-stimulatory conditions,^48^ or at a lower threshold glucose level.^48^ In contrast, a large number of cells (64%, Fig. 2) were metabolically uncoupled and largely unaffected by glucose flux, suggesting a specialized role distinct from glucose-dependent insulin release. Indeed, many groups have reported a high percentage of isolated primary β-cells that fail to respond to glucose.^4,44,46,49^ While this observation may initially seem at odds with a physiological role for these non-responding cells, if one considers recent research on primary β-cell electrical heterogeneity,^5,50,51^ within a functional gap-junction coupled electrical syncytium, then such an observation may be expected as follows. It is now known that there is a smaller subset of β-cells that respond rapidly to an increase in glucose and have been termed “first-responder” cells.^2,6^ There is also another larger population of “follower” β-cells that have low glucose responsiveness, and their excitability is controlled through a gap-junctional coupled multicellular network connected to the glucose-responsive cells. Interestingly, our modelling aligns well with the above concept as we observe a small population of glucose-responsive cells and a larger population of non-responding β-cells that may correspond to the “first responder” and “follower” β-cell phenotypes, respectively, that are observed in primary islet studies.

Previous models constructed with only right-shifted *I*_*Na*_ saw higher excitability with a larger dynamic range of glucose-responsive β-cells. ^18,32^ Our inclusion of left-shifted *I*_*Na,low*_ β-cells (63% of all *I*_*Na*_ in our optimized model), resulted in lower excitability than past rodent models, yet could not fully explain the decreased glucose responsiveness. We propose that other inhibitory components, like g_BK_, also play a substantial role in glucose excitability via modulation of resting membrane potential. Our estimate of variability for g_BK_ was taken from the foundation patch-clamp dataset for the Riz model,^32^ and reflects the iberiotoxin-sensitive current variability at +30 mV among n = 9 cells. Given the relatively small sample, it is certainly possible that this variability is an overestimate, but the same could be said for other components. Perhaps the most important conclusion to draw from this is that extending the fitting process to include larger patch-clamp datasets for human K^+^ currents could be beneficial. At least one such dataset does exist, and its inclusion is a priority for follow-up studies.^52^

### Importance for integrated function of the islet

A central finding of this study is that the bimodal heterogeneity of *I*_*Na*_ voltage-dependence is among the most important cellular factors influencing the proportion of human β-cells that are electrically active at basal glucose, and the proportion that remain silent in response to increased glucose. While these populations may seem secondary to cells that exhibit classical glucose-induced bursting, it is well known that dispersed primary β-cells rarely exhibit that classical phenotype, even though it is the dominant form of activity in intact islets.^53^ Instead, bursting of the islet network is thought to arise from functional heterogeneity among β-cells and the coupling between cells within the network.^53^ In this context, changes in the balance of basally active and glucose non-responsive cells are crucial to determining glucose-induced activity of the intact islet. To this point, *I*_*Na*_ has been assumed to be a lesser player in the activation of pancreatic islets in all species. These findings suggest that expression of NaV isoforms with differing voltage dependence offers a means of tuning overall islet excitability and dynamic responses to glucose.

### Challenges in studying β-cell populations

A final takeaway from our study is the necessity of large-scale modelling simulations to dissect the effects of cellular heterogeneity. Components like ḡ_KATP_, GK, and g_BK_ retained detectable influence in smaller virtual sample sizes, but subtle influence from factors like *I*_*Na*_ and *I*_*Ca*_ required robust sampling. For example, *I*_*Na*_ voltage dependence plays a substantial role in dictating baseline excitability but would have likely been overlooked without our large simulated population of 3,000 cells. These findings emphasize the understanding that subtle forms of intercellular heterogeneity require a scalable model such as our PoM simulations to assess physiological relevance.

### Limitations and future directions

We have noted several study limitations, which have motivated our continuous model refinement. Firstly, the Camunas-Soler *et al*. patch-clamp dataset was used for fitting most parameters focused on inward current. As such, the heterogeneity of outward conductances (ḡ_KATP_, g_BK,_ g_Kv_) was inferred from smaller and potentially less representative datasets. Incorporating newer, large-scale voltage-clamp datasets will help us refine the model sensitivity and fidelity, particularly in K+ current species.

Next, our implementation of bimodal *I*_*Na*_ inactivation relied on fixed proportions rather than clustering from single-cell data. This likely underestimates the full impact of *I*_*Na*_ density, providing a conservative estimate of its contribution to shaping network excitability. Future work must integrate covariations between NaV isoform expression, window current, and other electro-metabolic properties to reflect a more realistic electrophysiological phenotype.

Finally, our simulations were conducted in uncoupled β-cells rather than networked cells. Intact islets exhibit a steeper glucose activation curve and a higher threshold for excitability, properties driven by functional coupling and network coordination as discussed above.^45,46,49,54–56^ While the directionality of our simulations is clear, their physiological magnitude and relevance to whole-islet function must be validated in multicellular computational models. Such models must include gap junctional coupling, paracrine signalling, and non-β-cells to maximize physiological realism.

## CONCLUSION

PoM simulations offer new insight into how cell-intrinsic electro-metabolic heterogeneity contributes to human β-cell excitability and glucose responsiveness and establish a generalizable modelling framework to further interrogate cellular heterogeneity. Our model suggests that bimodal *I*_*Na*_ voltage dependence and channel kinetics are a critical determinant of basal excitation, glucose metabolism selectively induces electrical bursting, and human β-cells have weaker electro-metabolic coupling than rodent β-cells. Such observations were enabled through large-scale modelling informed by high-quality primary human data. Without hypothesis-independent primary data collection and dataset generation, capturing physiologically relevant β-cell heterogeneity would not be possible. As the resolution and breadth of single-cell datasets continue to improve, data-driven computational models will be essential for interpreting functional variability and informing testable hypotheses.

## ACKNOWLEDGEMENTS

We would like to acknowledge the contribution of Karoline Horgmo-Jæger for sharing her excellent initial implementation of the baseline Riz *et al*. model with us. This research was funded by a Canadian Institutes of Health Research (CIHR) Project grant (P.E.L) and CIHR Doctoral student award (J.K). P.E.M holds the Canada Research Chair in Islet Biology.

## METHODS

### Human β-cell model

#### Baseline model

Using the human β-cell model published by Riz *et al*.^18^, we established a baseline β-cell for our initial simulations. This model captures several key elements of glucose metabolism, including four reactions catalyzed by glucokinase (GK), phosphofructokinase (PFK), fructose-1,6-bisphosphate aldolase (FBA), and glyceraldehyde-3-phosphate dehydrogenase (GAPDH). Oscillatory behaviour is driven by positive feedback of fructose-1,6-biphosphate (FBP) on PFK.^39^ ATP production is abstracted into a mimetic variable, *a*, which mediates electro-metabolic coupling by approximating expected changes in ATP availability and modulating ATP-sensitive K+ current (*I*_*KATP*_). The dynamics of *a* are modelled as the balance of terminal glycolytic flux *V*_*GAPDH*_ and the ATP consumption coefficient *kA*:

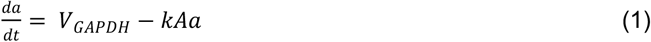

This variable, in turn, suppresses the fractional activation of *I*_*KATP*_ via:

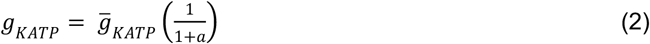

### Bimodal I_Na_ implementation

To capture the bimodal voltage dependence of *I*_*Na*_ observed by Camunas-Soler *et al*.^36^, we implemented two distinct sodium channel variants: *I*_*Na,high*_, with more depolarized steady-state activation and inactivation voltages, and *I*_*Na,low*_ with more hyperpolarized voltage-dependence. Although our model remains agonistic to specific isoforms, the literature suggests that SCN9a (Nav1.7), SCN8a (Nav1.6) and SCN3a (Nav1.3) are present in human β-cells.

Each β-cells in the population was assigned only one form of *I*_*Na*_, controlled by the population-level parameter *ρ. ρ* represents the fraction of cells expressing *I*_*Na,high*_ with the remainder of cells (1-*ρ*) expressing *I*_*Na,low*_. Both variants follow the same instantaneous activation kinetics with rapid inactivation (*τ*_*h*_ = 2 ms). Mean parameter values are summarized in Supplementary Table 1.^18^

### Modelling heterogeneity via Population of Models

We employed a hybrid method for implementing heterogeneity across simulated β-cells. Two primary approaches exist in previous literature: (1) parameter variation based on reported sample variance without fitting,^6,11,12,14,57,58^ and (2) fitting heterogeneous parameter distributions to functional data through data trimming.^28,29,51,59–61^ To avoid distortion of parameter distributions, we did not implement data trimming or unconstrained sampling. Instead, we applied differential evolution (DE) optimization to fit coefficients of variation (*σ*) for selected parameters to match experimentally derived distributions.^36^ All parameter variations followed lognormal distribution to ensure physiological plausibility and appropriate fold-symmetry.

#### Experimental dataset and reference data

We used electrophysiological patch-clamp recordings from 180 human β-cells published by Camunas-Soler *et al*.^36^ These cells were subjected to standardized voltage-clamp protocols at 1mM or 10mM extracellular glucose. Since these protocols eliminated the K^+^ current, it is unlikely that glucose contributed significantly to the variation in measured inward currents. Because the Baseline model does not include synaptic fusion and associated changes in capacitance, we used changes in intracellular [Ca^2+^] as a proxy for exocytosis. Similar to analysis of the experimental capacitance responses to the pulse-train protocol, we used the peak change in [Ca^2+^] during the first pulse to indicate early exocytosis and the peak change (from pre-protocol baseline) during the final pulse to indicate total exocytosis.

#### Heterogeneous parameters

Heterogeneity was implemented as a coefficient of variation in a subset of 16 parameters: 14 electrophysiologic and 2 metabolic (Table 1). For five electrical parameters, variability was derived from smaller studies with higher resolution but lower sample sizes. The remaining nine electrophysiologic parameters were fitted to voltage-clamp distributions published by Camunas-Soler *et al*. ^36^ For metabolic parameters (maximal GK flux, *V*_*GK,max*_, and the glucose concentration at half-maximal GK binding *K*_*GK*_), we used inter-patient variability as a conservative estimate for intercellular heterogeneity.

Similar to prior studies, we construct the population from the baseline model by defining a matrix of scaling factors *S*, where s_*i,j*_ is the scaling factor for the *j*_th_ parameter of the *i*_th_ cell in the population. These scaling factors are simple coefficients for the baseline parameter vector 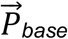, Supplemental Table 1), with a mean of 1 and a standard deviation of *σ*. To convert from the scaling factor matrix to the parameter vector 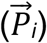 of the *i*_th_ cell is simply the elementwise (Hadamard) product:

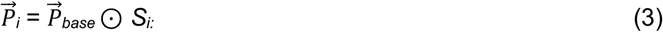

Table 1 lists both initial estimates of *σ* for all varied parameters (from prior studies) and final *σ* values for parameters optimized by DE. Parameters for which human-specific variation was not available were left at their default value in the baseline model. That is, for these *k* invariant parameters, *σ*_*k*_ = 0 and *S*_:,*k*_ = 1.

#### Parameter sampling

Using *σ* values (initial estimates or DE-optimized; see Table 1), we constructed S by lognormal sampling for 3,000 cells. This lognormal sampling method ensured that scaling factors were positive values along a symmetric log scale.^16,29,31,62^ The resultant parameters were skewed towards larger absolute values, where the skew increases with *σ*. Importantly, these parameters reflect the variance observed in the Camunas-Soler *et al*. dataset despite not being a feature of conventional Gaussian sampling. Additional constraints were applied to *I*_*Na*_ voltage dependence parameters to avoid physiologically impossible window currents and slope factors. These constraints are detailed in the supplementary information.

#### Fitting parameter variability to voltage clamp behaviour

We fit parameters 6-17 from Table 1 using DE to deliberately minimize deviations between model output and the primary human β-cells voltage clamp dataset. While divergent from previous modelling approaches,^1,11–14,16,57,58^ this method allowed us to simultaneously capture contributions from overlapping ion channel species (*I*_*Ca*_ and *I*_*Na*_)and avoid *a priori* assumptions about which parameters drive specific metrics. Sensitivity analyses confirmed the cross-contributions between each current type (Figure S2). ^61^ The DE process also optimized the population level *ρ* for *I*_*Na*_ . Full optimization details are provided in the Supplementary Information.

### Simulating glucose responses

The final population of 3,000 β-cells was simulated for 600 seconds at 2 (low) and 20 mM (high) glucose. Voltage traces were classified into four electrical phenotypes: Silent, Bursting, Spiking, or Depolarized. The classification was based on a modified version of Andrean *et al*.’s algorithm (see Figure S5). ^16^ We next grouped cell types by their transitional phenotype across glucose concentrations and analyzed ATP-mimetic values to define metabolic activation. A threshold of *a* = 2.0mM was used to classify simulations as metabolically active.

### Population-level optimization of electro-metabolic coupling

Since glucose responsiveness was not a constraint in our initial parameter fitting, we assessed whether emergent β-cell activity matched expectations from primary data. Rodent studies suggest a dynamic range of ∼50% electrical activation upon glucose stimulation, whereas our default model showed lower fidelity (19% to 35%; dynamic range of 16%). To increase our dynamic range, we adjusted the ATP consumption coefficient *kA* (Equation 1) to 4e^-05. With this value, the fraction of electrically active cells increased to 49% at 20 mM glucose, with low glucose activity unchanged (22%), yielding an improved coupling fidelity of 0.47. Lowering *kA* further increased activation but disrupted the Bursting phenotype. All analyses of “sensitized” populations use *kA* = 4e^-05

### Statistical analysis

Our statistical analyses ranked the phenotypic influence of model parameters within each glucose level, as well as for phenotype transitions from 2 to 20 mM glucose. We took two approaches. First, conventional group-wise tests (e.g. ANOVA, Kruskal-Wallis) to identify mean (or median) differences. These analyses were chosen to be analogous to experimental procedures and assume independence among the parameters. Second, we applied logistic regression models to binarized phenotypes and z-scored parameter deviations, which allowed us to rank parameters by their influence without assuming independence or equal variance. This partitions simultaneous variation among the parameters and returns weights that indicate the sensitivity of the phenotype to the parameters. Full details are provided in the Supplementary Information.

## Code availability

All code will be made available with corresponding experimental data at: https://github.com/andygedwards/BCell-Populations

