## Supplementary material for "Computational modelling of natural cell-to-cell heterogeneity reveals key parameters that control the diversity of human pancreatic islet β-cell excitability in response to glucose"

### Supplemental Information

#### Baseline Riz *et al.* β-cell model.

All parameters for the baseline Riz *et al.*[18] model are provided in Supplemental Table 1. These include parameters for both components of $I_{Na}$ ($I_{Na,high}$ and $I_{Na,low}$). Each component has identical form to the original representation of $I_{Na}$ in Riz *et al.,* and the default $I_{Na,high}$ parameters are identical to those of the original $I_{Na}$.

| Parameter Name | Parameter Value | Parameter Units | Parameter Description |
|---|---|---|---|
| **High-voltage inactivated sodium current ($I_{Na,high}$)** | | | |
| $V_{hNa,high}$ | $-42.00$ | $mV$ | $I_{Na,high}$ half inactivation potential |
| $V_{mNa,high}$ | $-18.00$ | $mV$ | $I_{Na,high}$ half activation potential |
| $g_{Na,high}$ | $0.40$ | $nS/pF$ | $I_{Na,high}$ maximal conductance |
| $n_{hNa,high}$ | $6.00$ | $mV$ | $I_{Na,high}$ inactivation slope factor |
| $n_{mNa,high}$ | $-5.00$ | $mV$ | $I_{Na,high}$ activation slope factor |
| $\tau_{hNa,high}$ | $2.00$ | $ms$ | $I_{Na,high}$ inactivation time constant |
| **Low-voltage inactivated sodium current ($I_{Na,low}$)** | | | |
| $V_{hNa,low}$ | $-71.00$ | $mV$ | $I_{Na,low}$ half inactivation potential |
| $V_{mNa,low}$ | $-43.00$ | $mV$ | $I_{Na,low}$ half activation potential |
| $g_{Na,low}$ | $0.40$ | $nS/pF$ | $I_{Na,low}$ maximal conductance |
| $n_{hNa,low}$ | $2.50$ | $mV$ | $I_{Na,low}$ inactivation slope factor |
| $n_{mNa,low}$ | $-5.00$ | $mV$ | $I_{Na,low}$ activation slope factor |
| $\tau_{hNa,low}$ | $2.00$ | $ms$ | $I_{Na,low}$ inactivation time constant |
| **L-type $Ca^{2+}$ current ($I_{CaL}$)** | | | |
| $V_{mCaL}$ | $-25.00$ | $mV$ | $I_{CaL}$ half activation potential |
| $g_{CaL}$ | $0.14$ | $nS/pF$ | $I_{CaL}$ maximal conductance |
| $n_{mCaL}$ | $-6.00$ | $mV$ | $I_{CaL}$ activation slope factor |
| $\tau_{hCaL}$ | $20.00$ | $ms$ | $I_{CaL}$ activation time constant |
| **PQ-type $Ca^{2+}$ current ($I_{CaPQ}$)** | | | |
| $V_{mCaPQ}$ | $-10.00$ | $mV$ | $I_{CaPQ}$ half activation potential |
| $g_{CaPQ}$ | $0.17$ | $nS/pF$ | $I_{CaPQ}$ maximal conductance |
| $n_{mCaPQ}$ | $-6.00$ | $mV$ | $I_{CaPQ}$ activation slope factor |
| **T-type $Ca^{2+}$ current ($I_{CaT}$)** | | | |
| $V_{hCaT}$ | $-64.00$ | $mV$ | $I_{CaT}$ half inactivation potential |
| $V_{mCaT}$ | $-40.00$ | $mV$ | $I_{CaT}$ half activation potential |
| $g_{CaT}$ | $0.05$ | $nS/pF$ | $I_{CaT}$ maximal conductance |
| $n_{hCaT}$ | $8.00$ | $mV$ | $I_{CaT}$ inactivation slope factor |
| $n_{mCaT}$ | $-4.00$ | $mV$ | $I_{CaT}$ activation slope factor |
| $\tau_{hCaT}$ | $7.00$ | $ms$ | $I_{CaT}$ inactivation time constant |
| **Small conductance $Ca^{2+}$-activated $K^{+}$ current ($I_{SK}$)** | | | |
| $K_{SK}$ | $0.57$ | $\mu M$ | $I_{SK}$ half-maximal $Ca^{2+}$ concentration |
| $g_{SK}$ | $0.10$ | $nS/pF$ | $I_{SK}$ maximal conductance |
| $n_{SK}$ | $5.20$ | $\mu M$ | $I_{SK}$ Hill coefficient for $Ca^{2+}$ activation |
| **Big conductance $Ca^{2+}$-activated $K^{+}$ current ($I_{BK}$)** | | | |
| $B_{BK}$ | $20.00$ | $pA/pF$ | $I_{BK}$ $Ca^{2+}$-independent current |
| $V_{mBK}$ | $0.00$ | $mV$ | $I_{BK}$ half activation potential |
| $\bar{g}_{BK}$ | $0.02$ | $mS/\mu A$ | $I_{BK}$ maximal conductance |
| $n_{mBK}$ | $-10.00$ | $mV$ | $I_{BK}$ activation slope factor |
| $\tau_{mBK}$ | $2.00$ | $ms$ | $I_{BK}$ activation time constant |
| **Delayed rectifier $K^{+}$ current ($I_{Kv}$)** | | | |
| $V_{mKv}$ | $0.00$ | $mV$ | $I_{Kv}$ half activation potential |
| $g_{Kv}$ | $1.00$ | $nS/pF$ | $I_{Kv}$ maximal conductance |
| $n_{mKv}$ | $-10.00$ | $mV$ | $I_{Kv}$ activation slope factor |
| $\tau_{mKv0}$ | $2.00$ | $ms$ | $I_{Kv}$ activation time constant |
| **ATP-sensitive $K^{+}$ current ($I_{KATP}$)** | | | |
| $g_{KATP0}$ | $0.01$ | $nS/pF$ | Glycolysis disabled $I_{KATP}$ maximal conductance |
| $\bar{g}_{KATP}$ | $0.05$ | $nS/pF$ | $I_{KATP}$ maximal conductance |
| $glycolysis$ | $1.00$ | $binary$ | glycolysis flag |
| **$K^{+}$ leak current ($I_{leak}$)** | | | |
| $V_{leak}$ | $-30.00$ | $mV$ | $I_{leak}$ equilibrium potential |
| $g_{leak}$ | $0.015$ | $nS/pF$ | $I_{leak}$ maximal conductance |
| **$Ca^{2+}$ handling proteins** | | | |
| $J_{SERCA,max}$ | $0.06$ | $\mu M/ms$ | Maximal SERCA flux |
| $K_{SERCA}$ | $0.27$ | $\mu M$ | $Ca^{2+}$ concentration at half-maximal SERCA transport |
| $J_{PMCA,max}$ | $0.021$ | $\mu M/ms$ | Maximal PMCA flux |
| $K_{PMCA}$ | $0.50$ | $\mu M$ | $Ca^{2+}$ concentration at half-maximal PMCA transport |
| $J_{leak}$ | $9.4 \times 10^{-4}$ | $\mu M/ms$ | $Ca^{2+}$ leak flux from SR to cytosol |
| $J_{NCX0}$ | $1.87 \times 10^{-2}$ | $ms^{-1}$ | NCX-mediated $Ca^{2+}$ extrusion flux |
| **Metabolic parameters** | | | |
| $G$ | $10.00$ | $mM$ | Glucose concentration |
| $K_{FBA}$ | $5.0 \times 10^{-3}$ | $mM$ | Fructose 1,6-bisphosphate concentration at half-activation of FBA |
| $K_{GAPDH}$ | $5.0 \times 10^{-3}$ | $mM$ | Glyceraldehyde 3-phosphate concentration at half-activation of GAPDH |
| $K_{GK}$ | $8.00$ | $mM$ | Glucose concentration at half-activation of GK |
| $K_{GPI}$ | $0.30$ | $unitless$ | Equilibrium constant for glucose-6-phosphate isomerase reaction |
| $K_{PFK}$ | $4.00$ | $mM$ | Fructose 6-phosphate concentration at half-activation of FBA |
| $K_{TPI}$ | $4.55 \times 10^{-2}$ | $unitless$ | Equilibrium constant for triose-phosphate isomerase reaction |
| $P_{FBA}$ | $0.50$ | $mM$ | Glyceraldehyde 3-phosphate concentration at half-saturation of FBA reverse reaction |
| $Q_{FBA}$ | $0.275$ | $mM$ | Dihydroxyacetone phosphate concentration at half-saturation of FBA reverse reaction |
| $V_{FBA,max}$ | $1.39 \times 10^{-4}$ | $mM/ms$ | Maximum flux of FBA |
| $V_{GAPDH,max}$ | $1.4 \times 10^{-3}$ | $mM/ms$ | Maximum flux of GAPDH |
| $V_{GK,max}$ | $5.56 \times 10^{-5}$ | $mM/ms$ | Maximum flux of GK |
| $V_{PFK,max}$ | $5.56 \times 10^{-4}$ | $mM/ms$ | Maximum flux of PFK |
| $X_{PFK}$ | $0.01$ | $mM$ | Fructose 1,6-bisphosphate concentration at half-saturating allosteric activation of PFK |
| $\alpha_{G}$ | $5.00$ | $unitless$ | Maximal allosteric effect of Fructose 1,6-bisphosphate on PFK half-activation |
| $h_{GK}$ | $1.70$ | $unitless$ | Hill coefficient of GK |
| $h_{PFK}$ | $2.50$ | $unitless$ | Hill coefficient of PFK in the absence of Fructose 1,6-bisphosphate |
| $h_{X}$ | $2.50$ | $unitless$ | Hill coefficient of allosteric effect of Fructose 1,6-bisphosphate on PFK half-activation |
| $h_{act}$ | $1.00$ | $unitless$ | Hill coefficient of PFK at saturating Fructose 1,6-bisphosphate concentration |
| $k_{A}$ | $1.0 \times 10^{-4}$ | $ms^{-1}$ | Pseudo-ATP degradation rate |
| **Buffering and diffusion parameters** | | | |
| $B$ | $0.10$ | $ms^{-1}$ | Diffusive flux constant from submembrane space to cytosolic space |
| $Cm$ | $10.00$ | $pF$ | Membrane capacitance |
| $Vol_{c}$ | $1.15 \times 10^{-12}$ | $L$ | Cytosolic volume |
| $Vol_{m}$ | $1.00 \times 10^{-13}$ | $L$ | Submembrane volume |
| $\alpha$ | $5.18 \times 10^{-15}$ | $\mu mol/pA/ms$ | Faraday-based constant to convert total $Ca^{2+}$ current to flux |
| $f$ | $0.01$ | $unitless$ | Fraction of $Ca^{2+}$ remaining free in cytosol |

**Supplemental Table 1: Baseline model parameters**

Spike repolarization and frequency dynamics in the baseline model are dominated by 3 activity-dependent $K^+$ currents. They are an ultra-rapidly activating and non-inactivating delayed rectifier current ($I_{Kv}$), and both small- ($I_{SK}$) and big-conductance ($I_{BK}$) $Ca^{2+}$-activated $K^+$ currents. $Ca^{2+}$ influx is mediated by T-, L-, and PQ-type $Ca^{2+}$ channels, and is returned to equilibrium by activity of the plasma membrane $Ca^{2+}$-ATPase (PMCA), sodium-calcium exchanger (NCX) and sarco-endoplasmic reticulum $Ca^{2+}$-ATPase (SERCA). $Ca^{2+}$-dependent exocytosis and secretory cascades are not implemented in the model and thus $[Ca^{2+}]$ serves as the sole functional proxy for insulin release. Of note, we apply the default configuration of the original model, which ignores the influence of the human ether a-go-go related gene (hERG) $K^+$ channel, and gamma-aminobutyric acid receptor activity. Additionally, we note that this model does not include the influence of local $Ca^{2+}$ nanodomains on $Ca^{2+}$-dependent $K^+$ channel function.[63] This property is thought to result from SR and plasma membrane channel co-localization, and while those dynamics may be important for some oscillatory behaviors in the islet, they are not incorporated here. The full parameter set for all components of the baseline model is provided in Supplemental Table 1.

**Human β-cell dataset.**

The dataset applied here is a reduction of the publicly available patch-seq dataset published by Camunas-Soler *et al*.[36] We performed our analyses on all transcript-confirmed β-cells, from healthy donors, which had not been cryopreserved, but had been patch-clamped. All β-cells fitting these constraints and which had non-zero values for at least one of the 6 electrophysiological metrics were included. This number amounted to 180 cells. All relevant datasets and code for reduction are provided in the code repository associated with this study.

**Structure of the scaling factor matrix and parameter sampling.**

We construct the scaling factor matrix ($S$) as mentioned in the main text. This matrix is rectangular, $\boldsymbol{n}$ (cells) x 16 (varied parameters). For each row (cell, $i$), the 16 columns (scaling factors, $j$) define a variation to the baseline model. The parameter $\rho_{high}$ operates at the population level to define the fraction of cells expressing $I_{Na,high}$. Therefore, scaling factors for $I_{Na,high}$ were included in $\rho_{high}\bullet\boldsymbol{n}$ rows and the $I_{Na,low}$ columns for those rows were set to zero, thus eliminating any contribution from $I_{Na,low}$ in those cells. The reciprocal process was applied to the remaining $(1\text{-}\rho_{high})\bullet\boldsymbol{n}$ rows, which included scaling factors for $I_{Na,low}$, but not $I_{Na,high}$.

*Sampling non-voltage dependent parameters:* Lognormal sampling was carried out independently for each parameter except the $I_{Na}$ voltage dependence parameters, which were varied in coordination to avoid unphysiologic window currents and slope factors. To be explicit, we did not assume *a priori* covariation among any of the parameters aside from those involved in $I_{Na}$ voltage dependence. While it is very likely that genuine biologic covariation occurs at the level of transcriptional control for many parameters, for example channel conductances ($g_{Na}$, $g_{CaL}$ etc.),[64–67] the low transcript number of most ion channels make data constraints for these covariances highly uncertain.

To perform the sampling, the mean ($\mu_N$) and standard deviation ($\sigma_N$) of the normal distribution underlying the final log-transformed scaling factor distributions were calculated as:

$$\mu_N = \ln\left(\frac{\mu_{est}^2}{\sqrt{\sigma_{est} + \mu_{est}^2}}\right)$$

and:

$$\sigma_N = \sqrt{\ln\left(\frac{\sigma_{est}}{\mu_{est}^2} + 1\right)}$$

Where $\mu_{est} = 1$ is the mean scaling factor estimate for all parameters (i.e. equivalent to default parameter values in the baseline model, Table S1), and the $\sigma_{est}$ values are shown in Table 1 as both the initial estimates from data sources, and the final set of $\sigma_{est}$ values after optimization.

<u>*Sampling $I_{Na}$ voltage-dependence parameters*</u>: For the $I_{Na}$ activation and inactivation parameters ($V_{hNa}$, $V_{mNa}$, and $n_{hNa}$), we applied 2 additional constraints. First, for both the $I_{Na,high}$ and $I_{Na,low}$ forms of the current, we varied these three parameters in coordination. That is, for each cell in the population the scaling factors for $V_{hNa}$ and $V_{mNa}$ were identical, and sampled from a lognormal distribution. Because the experimental inactivation v-half distributions for both forms of $I_{Na}$ skew toward zero, rather than away, we also shifted the scaling factor skew to match this direction. Formally, if $\vec{s}$ is a vector of scaling factors that have been randomly sampled from a lognormal distribution (calculated from $\mu_{est} = 1$, and $\sigma_{est}$ for either $V_{hNa}$ or $V_{mNa}$), the final scaling factor vectors for the v-half of inactivation ($\vec{s}_{Vh}$) and activation ($\vec{s}_{Vm}$) are:

$$\vec{s}_{Vh} = \vec{s}_{Vm} = 2\mu_{est} - \vec{s}$$

The scaling factor vector ($\vec{s}_{nh}$) for the inactivation slope factor ($n_{hNa}$) was then calculated from the same $\vec{s}$ as:

$$\vec{s}_{nh} = (\vec{s} + 0.5)/1.5$$

These sampling procedures were performed identically but independently for $I_{Na,high}$ and $I_{Na,low.}$

We applied these constraints because preliminary simulations (without the constraints) often resulted in very large window currents, or unphysiologic inactivation slope factors. The result for heterogeneity of these parameters is that their variations are perfectly correlated to each other (negatively correlated for $n_{hNa}$). While we are unaware of data that specifically describe coordinated variation in activation and inactivation voltage-dependence in $I_{Na}$, this remains a necessary assumption of our approach. It is also clear that discoordination of these processes underlies major $I_{Na}$ channelopathies, such as type 3 long QT syndrome (LQT3).[68]

The final scaling factor distributions for each parameter are shown in Figure S1, where the degree of variation in each parameter is reflected in the relative skew of the distribution. This figure also shows the reversed skew direction (i.e. slightly towards 0) and coordinated variation for the $I_{Na}$ voltage-dependence parameters.

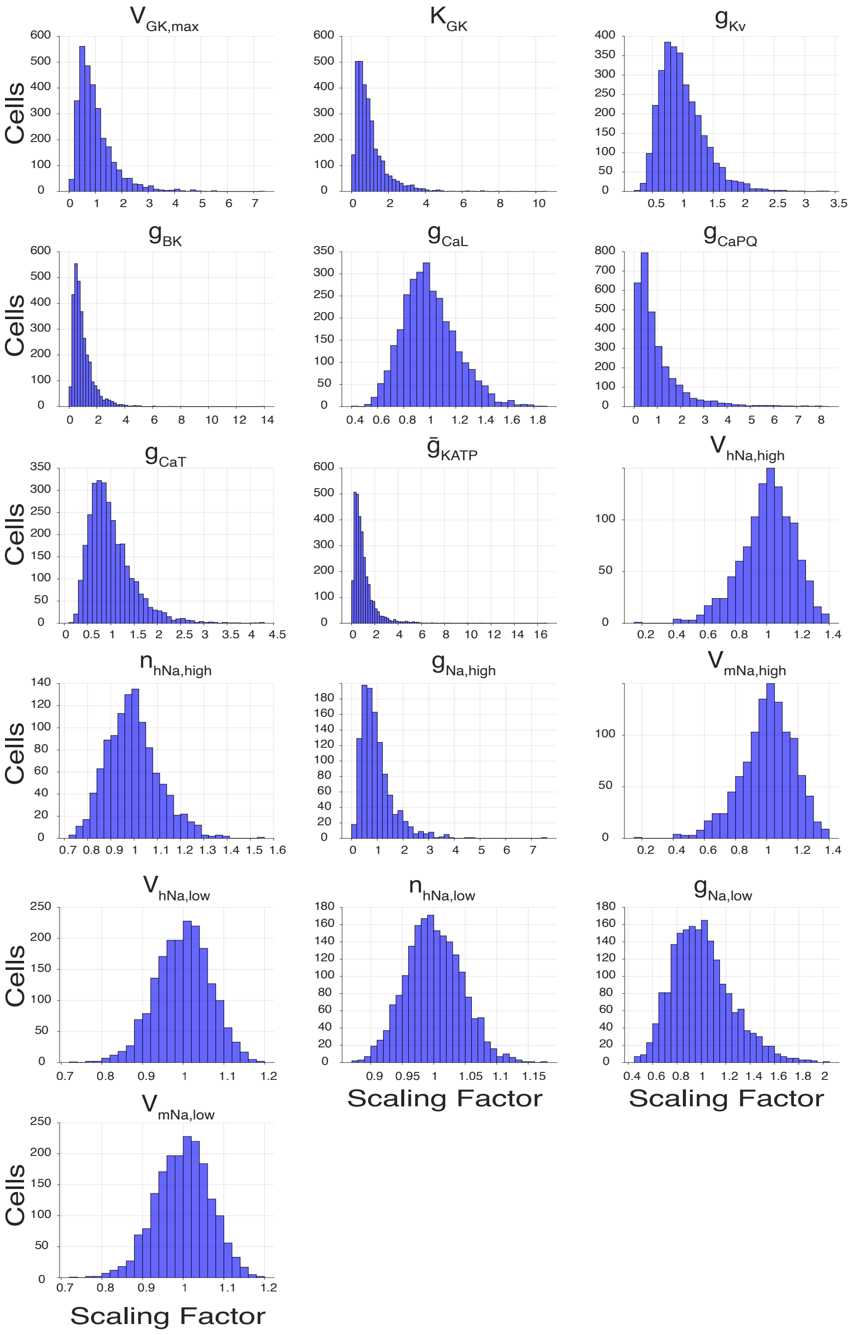


**Figure S1. Lognormal distributions of the heterogeneous parameters after optimization.**

#### Sensitivity Analyses.

As mentioned in the main text, the compound voltage-clamp protocols used here and in Camunas-Soler et al.,[36] are likely to involve some degree of cross-contribution between $I_{Na}$ and $I_{Ca}$ in several of the measured metrics. To quantify that cross-contribution and to provide better understanding of how the variable scaling factors effect the voltage-clamp metrics, we performed a global sensitivity analysis using partial least squares regression for each metric.[59,61] The regression coefficients from these analyses provide a linear approximation of each scaling factor's weighting for determining the metric of interest, and are visualized as bars in Figure S2. Each bar indicates the sensitivity (of the metric with respect to changes in the scaling factor, and are in terms of the absolute values of each. Thus, negative sensitivities indicate that the metric absolute value is inversely related to the scaling factor, and the magnitude indicates the sensitivity of the relationship. Because the variability of each parameter also differed, it is important to consider that range (Fig S1) to interpret the influence of that parameter on the metric. It is clear from these analyses that $I_{Ca}$ parameters do have some influence on metrics of $I_{Na}$ and vice versa. However, when relative variation in each parameter is also taken into account, the separation is reasonably specific for the influence of $I_{Ca}$ parameters on the two $I_{Ca}$ metrics. That specificity increases for the role of $I_{Na}$ parameters on the $I_{Na}$ metrics, and is quite specific for the influence of $I_{Ca}$ parameters (particularly $g_{CaPQ}$) on the exocytotic metrics. Because voltage is clamped and K+ currents are not evoked in these protocols, we see essentially no influence of either metabolic or $K^+$ current parameters on any metric.

It should be noted that the cross-contribution of $I_{Ca}$ and $I_{Na}$ shown in Fig S2 is for the population after optimization by Differential Evolution. However, we noticed very similar sensitivities in all populations we tested. Thus, it is very likely that these are intrinsic properties of the model, and are also present to some degree in the experimental data set.

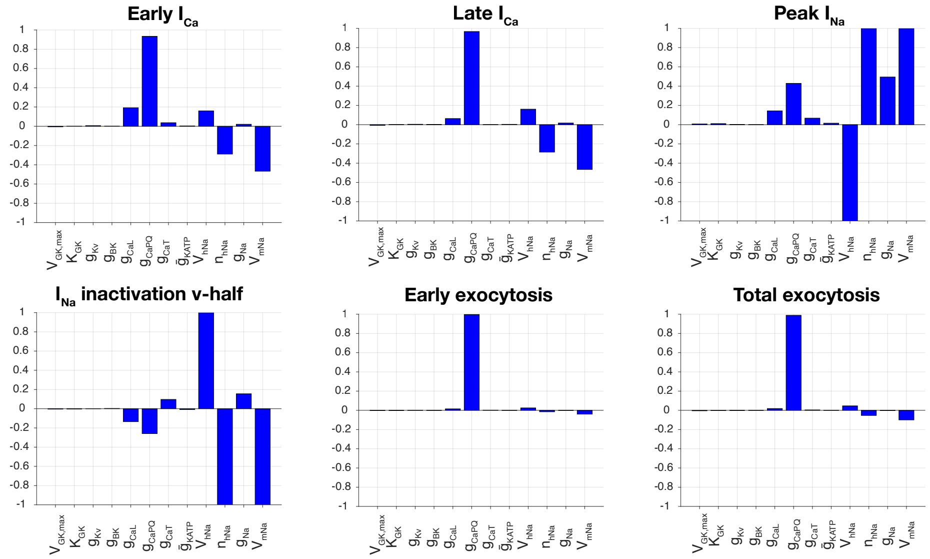


**Figure S2. Partial least squares global sensitivity analyses for the voltage-clamp metrics in the optimized population.**

#### Differential Evolution optimization of parameter heterogeneity.

The cross-contribution between $I_{Ca}$ and $I_{Na}$ shown above was our primary motivation for applying combinatorial optimization to define the heterogeneity of the $I_{Ca}$ and $I_{Na}$ parameters. An alternative would have been to directly take variations in $g_{Na}$ and $g_{Ca}$ calculated from the experimental dataset and assign fractions of each to the subtypes of $I_{Ca}$ (L, T, and PQ) and $I_{Na}$ (high and low) present in the model. This approach has been used by one previous study.[16] However, because it is unclear whether those fractions should be constant, and because it is clear that both $I_{Na}$ and $I_{Ca}$ are contributing to both sets of metrics in the model (and most likely the experimental dataset) we chose the optimization approach to avoid bias and invoke fewer assumptions.

We used several algorithms for computationally intensive objective functions in our preliminary tests (e.g. Genetic Algorithms, Particle Swarm Optimization), but Differential Evolution[69] consistently offered the best optimization trajectory and optimization error.

Each DE reconfiguration step involved varying the $\sigma$ values for parameters 6-16 (Table 1), and $\rho_{high}$, which were then applied to generate a new population of cells subjected to the 3 voltage clamp protocols assessing $I_{Na}$, $I_{Ca}$, and exocytosis. The resulting time-series were analysed as for the experimental time-series. The cost function for reconfiguration was calculated as a weighted multi-objective error between the model and experimental distributions for the v-half of $I_{Na}$ inactivation, peak $I_{Na}$, early $I_{Ca}$, late $I_{Ca}$, and early and total exocytosis. We used early $Ca^{2+}$ transient amplitude and total $Ca^{2+}$ amplitude as proxies for early and total exocytosis as capacitive changes resulting from exocytosis are not included in the Riz model.[18] The range within which the 10 $\sigma$*'s* were allowed to vary was ± 2- or 3-fold of their initial estimates, and depended on the parameter (see Table 1). Those initial estimates were taken from previous experimental studies which are also provided in Table 1 (Source). The fraction of cells expressing $I_{Na,high}$ has not been reported in humans, thus $\rho_{high}$ was allowed to vary between 0.1 and 1.0.

Because each optimization reconfiguration was references against the 180 experimental β cells, we constructed populations of only 200 cell models (at each parameter reconfiguration step. The final implementation of the DE employed 750 parameter reconfigurations within each “evolutionary population”, and we required at minimum 5 different evolutionary populations to be evaluated before convergence. Thus, at minimum a total of 3750 different 200 cell populations were submitted to the cost function for error calculation. Convergence criteria were met if the minimum cost in next evolutionary population did not improve by more than 3%. To verify the optimization results we reinitialized it at the optimal solution and ran a refined optimization by expanding the number of models from 200 to 1000, using another 2 evolutionary populations (1500 different 100- cell populations). The parameter-cost relationships for the 3 most determinant parameters ($g_{CaPQ}$, $\rho$, and $V_{hNa,low}$) for this refined optimization are shown in Fig S4. Lastly, we performed all glucose response analyses on both the optimal population (Figures 2 – 5), and the 2 populations with the next lowest cost function errors. The reported results were consistent across all of these populations.

The population shown in Figure 1 represents the best fit when all of the variable $\sigma$’s (parameters 6-17, Table 1) were included in the optimization. This optimization approach was particularly sensitive to $\sigma_{CaPQ}$ (Figures S2 and S4), up to the upper limit of $\sigma_{CaPQ}$ = 1.18. All further analyses presented in the main manuscript are for the population generated using this optimal (and high) heterogeneity for $g_{CaPQ}$. However, because $\sigma_{CaPQ}$ is unique in its ability to improve the fit to the ensemble of voltage clamp data, we also considered that such high $g_{CaPQ}$ heterogeneity may represent overfitting. To assess the implications for population function, we performed the optimization after fixing $\sigma_{CaPQ}$, $\sigma_{CaT}$ and $\sigma_{CaL}$ at their initial estimates from earlier small sample studies (Table 1, initial $\sigma$), and only optimizing $\sigma$'s for $I_{Na}$ (parameters 9-17, Table 1). These $I_{Na}$-only optimizations made only subtle differences to both the voltage-clamp metrics (Fig S3) and glucose responses. Thus we reverted to the original less-constrained optimization results.

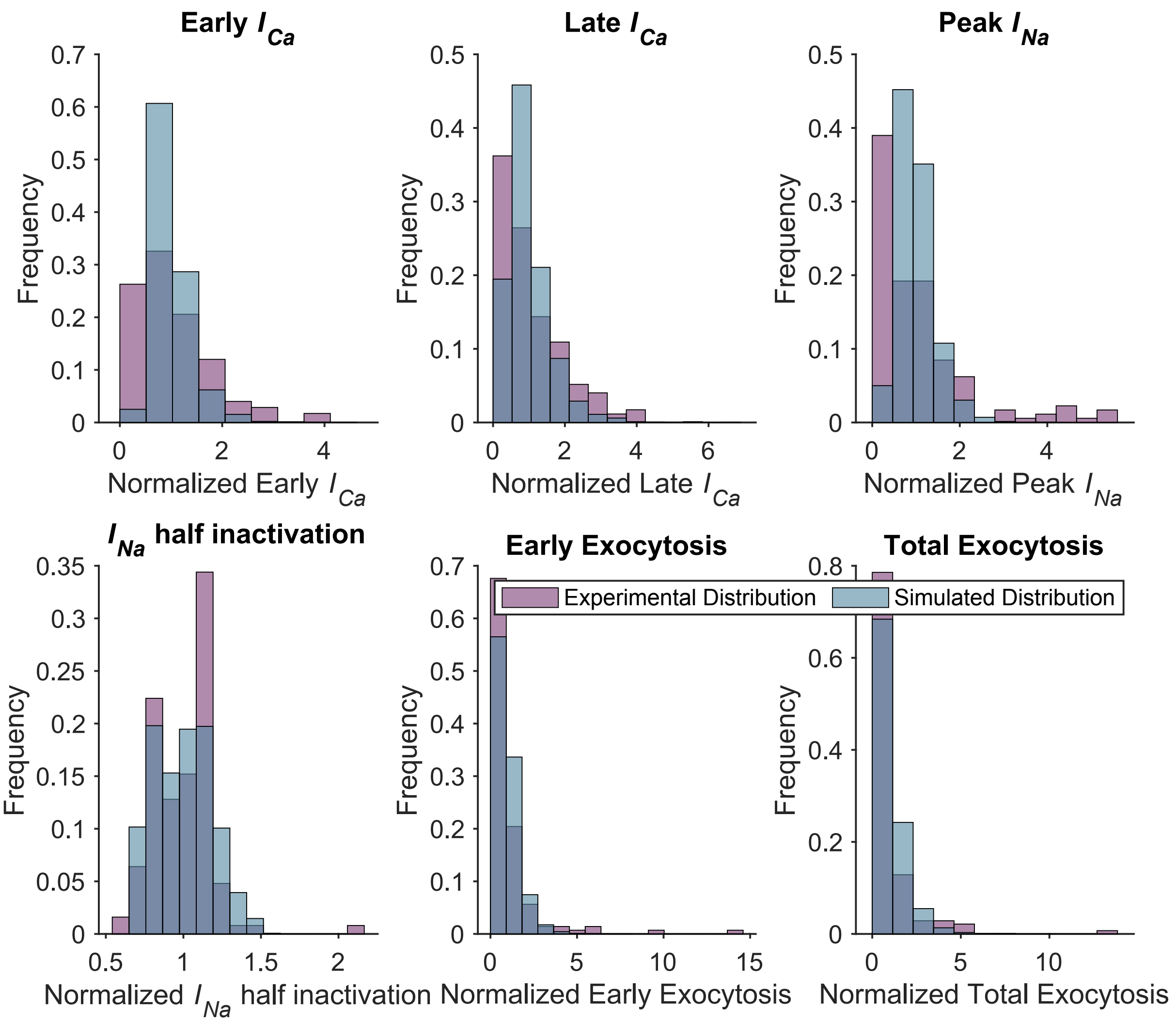


**Figure S3. Distributions for the voltage-clamp metrics in the population for which only $I_{Na}$ parameters were optimized. These exhibit very little difference to those returned by the full optimizations (Figure 1).**

*Cost function:*

The optimization error was defined by a multi-objective cost function. As mentioned above, the 200 models of the population were subjected to the experimental voltage protocols. Metrics for early $I_{Ca}$, late $I_{Ca}$, peak $I_{Na}$, $I_{Na}$ inactivation v-half, early exocytosis, and late exocytosis were extracted for each model.

Skewed and kurtotic Gaussian distributions were calculated for each resulting metric distribution, for both the 200 models and the 180 experimental cells. The distribution for $I_{Na}$ inactivation v-half was fitted to a double gaussian given the noted bimodality (Figure 1).

Each gaussian had 4 parameters for fitting ($\bar{x}$, σ, *s*, and *k*). $\bar{x}$ and $\sigma$ are the metrics mean and standard deviation, respectively, while, *s* and *k* are the skewness and kurtosis, which were calculated as:

Skewness: $s = \frac{E(x-\bar{x})^3}{\sigma^3}$, Where: $E(x - \bar{x})^3$ is the mean of the cubed deviations in the data, and $\sigma$ is the standard deviation.

Kurtosis: $k = \frac{E(x-\bar{x})^4}{\sigma^4}$, Where: $E(x - \bar{x})^4$ is the mean of the quartic deviations in the data, and $\sigma$ is the standard deviation.

To allow the experimental and simulated distributions of each metric to be directly compared across the same range of values we used kernel density estimation to fit probability density functions (PDF) for both. The error for each metric was then calculated as the RMS error between the simulated and experimental PDFs across that range.

Finally, the total multi-objective cost was the sum of these metric errors ($\epsilon$, n = 6) after each was multiplied by a corresponding weighting factor ($\theta$).

$$Total\ Cost = \sum_{i=1}^{n} \theta_i \epsilon_i$$

The $\theta$ weighting factors were applied to emphasize fitting of the $I_{Na}$ voltage dependent parameters, which were the most challenging to fit, and the final $\theta$ vector was:

1. Early exocytosis = 0.5
2. Total exocytosis = 0.5
3. Early $I_{Ca}$ = 1
4. Late $I_{Ca}$ = 1
5. Peak $I_{Na}$ = 1
6. $I_{Na}$ inactivation v-half = 3

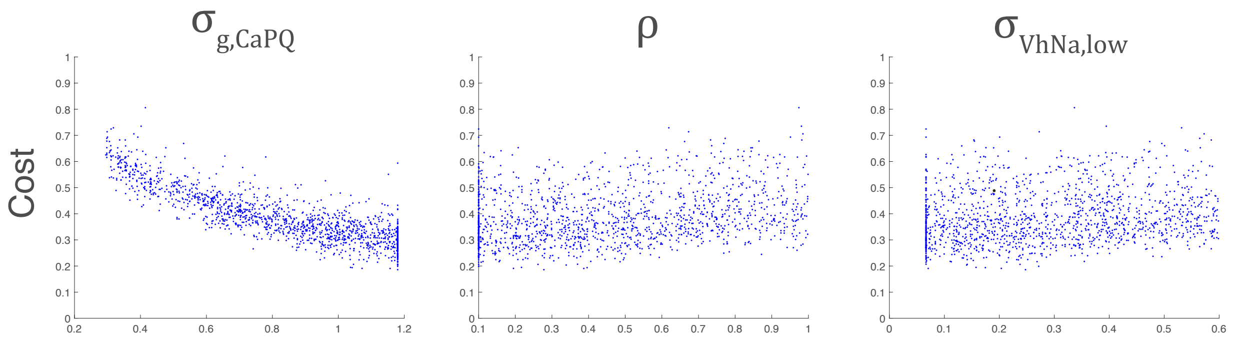


**Figure S4. Scatter plots showing the relationship between the 3 most determinant optimized parameters and the error returned by the multi-objective cost function. Note the relatively tight monotonic trajectory for $\sigma_{gCaPQ}$**

#### Glucose-response Classification.

All time series were classified by the algorithm shown in Fig S5. All series automatically classified as "Other" were manually classified as Silent, Spiking, or Depolarized. Greater than 1000 automatic classifications were randomly selected and verified for correctness by the same analyst. This algorithm is very similar to that developed by Andrean *et al.,* on a heterogeneous population built from the same dataset.[16]

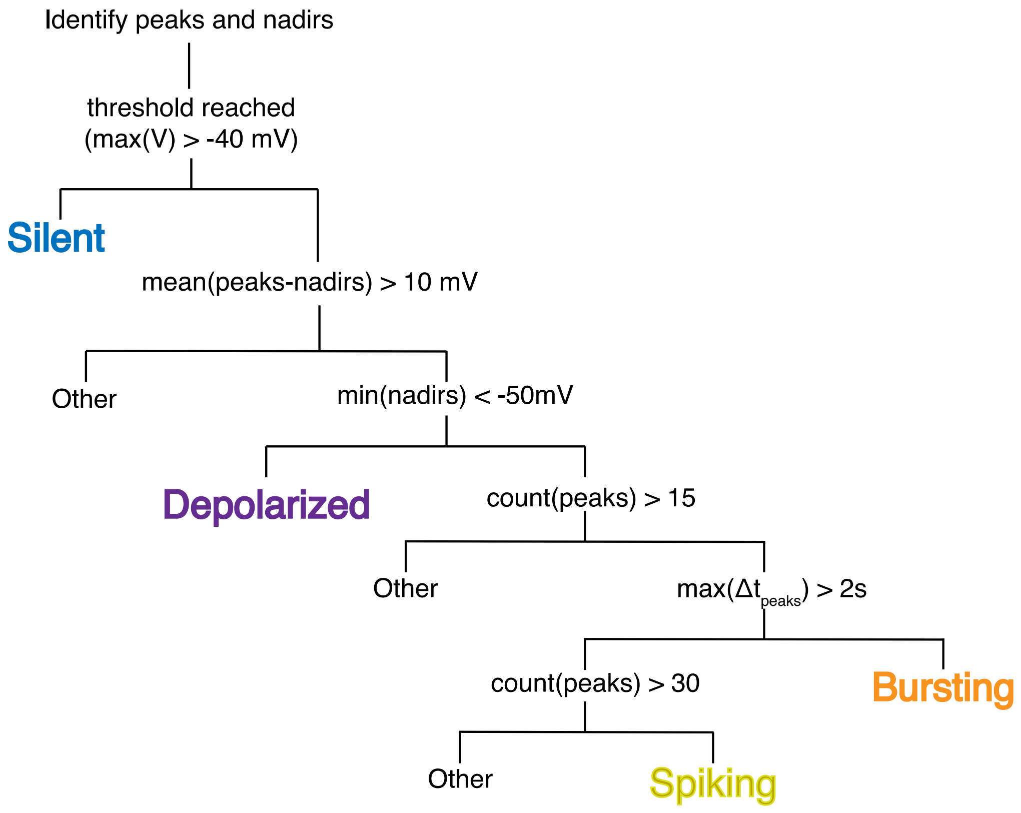


**Figure S5. Phenotype classification algorithm.**

**Statistical analyses of glucose-dependent phenotypes.**

Only Silent, Bursting, and Spiking cells were used for these analyses, as very few cells (< 1%) were classified as Depolarized or Other. As indicated in the main text, we approached these analyses using conventional group-wise (general linear and nonparametric) models, as well as logistic regression.

*Group-wise models:* Because the parameter distributions were not constructed as symmetric Gaussian (i.e. they were lognormally sampled), but may have become symmetric in their phenotype-specific groups, we first tested for normality (Andersen-Darling test) and homoscedasticity (Bartlett's test) among the 3 phenotype groups for each parameter. For normally distributed parameter groups we performed 1-factor ANOVAs (3 phenotype levels). For parameters that were not gaussian distributed we performed non-parametric Kruskal-Wallis omnibus tests. In each case, positive omnibus tests were followed by Dunn-Sidak post hoc tests for phenotype differences within each glucose concentration. Omnibus tests of glucose-dependent phenotype transitions were carried out similarly for each parameter, with the 3 levels corresponding to Silent-to-Silent, Silent-to-Bursting, and Silent-to-Spiking groupings. In that case, positive tests were followed by either the Dunnett (ANOVA omnibus) or Dunn (Kruskal-

Wallis omnibus) tests, with Silent-to-Silent treated as the reference (control) group. In all cases significance was defined at $\alpha = 0.05$.

*Logistic regression:* for each phenotype (Silent, Bursting, Spiking) or transition class (Silent-to-Silent, Silent-to-Bursting, Silent-to-Spiking), we dichotomized the classification to be positive or negative. We then performed logistic regression with the logit link function on standardized deviations of parameter values. The deviations were calculated as:

$$\frac{P_{ij} - \bar{P}_j}{\sigma_j}$$

$P_{ij}$ is the $j_{th}$ parameter for the $i_{th}$ cell, and $\bar{P}_j$ is the mean of the $j_{th}$ parameter for all cells. $\sigma_j$ is the standard deviation for $j_{th}$ parameter for all cells. This procedure returns Z-score deviations and allows the weights returned by logistic regression to be independent of their scales and therefore comparable as parameter sensitivities.

**Computational aspects.**

All simulations were performed with laptop scale compute power (Mac, M1, 8 cores, 16 GB memory). Code is currently implemented in Matlab (release 2023b), and upon publication will be made available with all source data to replicate major simulations and analyses at: https://github.com/andygedwards/BCell-Populations
